# A rapid and efficient screening system for neutralizing antibodies and its application for the discovery of potent neutralizing antibodies to SARS-CoV-2 S-RBD

**DOI:** 10.1101/2020.08.19.253369

**Authors:** Xiaojian Han, Yingming Wang, Shenglong Li, Chao Hu, Tingting Li, Chenjian Gu, Kai Wang, Meiying Shen, Jianwei Wang, Jie Hu, Ruixin Wu, Song Mu, Fang Gong, Qian Chen, Fengxia Gao, Jingjing Huang, Yingyi Long, Feiyang Luo, Shuyi Song, Shunhua Long, Yanan Hao, Luo Li, Yang Wu, Wei Xu, Xia Cai, Qingzhu Gao, Guiji Zhang, Changlong He, Kun Deng, Li Du, Yaru Nai, Wang Wang, Youhua Xie, Di Qu, Ailong Huang, Ni Tang, Aishun Jin

## Abstract

Neutralizing antibodies (Abs) have been considered as promising therapeutics for the prevention and treatment of pathogens. After the outbreak of COVID-19, potent neutralizing Abs to SARS-CoV-2 were promptly developed, and a few of those neutralizing Abs are being tested in clinical studies. However, there were few methodologies detailly reported on how to rapidly and efficiently generate neutralizing Abs of interest. Here, we present a strategically optimized method for precisive screening of neutralizing monoclonal antibodies (mAbs), which enabled us to identify SARS-CoV-2 receptor-binding domain (RBD) specific Abs within 4 days, followed by another 2 days for neutralization activity evaluation. By applying the screening system, we obtained 198 Abs against the RBD of SARS-CoV-2. Excitingly, we found that approximately 50% (96/198) of them were candidate neutralizing Abs in a preliminary screening of SARS-CoV-2 pseudovirus and 20 of these 96 neutralizing Abs were confirmed with high potency. Furthermore, 2 mAbs with the highest neutralizing potency were identified to block authentic SARS-CoV-2 with the half-maximal inhibitory concentration (IC50) at concentrations of 9.88 ng/ml and 11.13 ng/ml. In this report, we demonstrated that the optimized neutralizing Abs screening system is useful for the rapid and efficient discovery of potent neutralizing Abs against SARS-CoV-2. Our study provides a methodology for the generation of preventive and therapeutic antibody drugs for emerging infectious diseases.

## Introduction

Pandemic outbreaks of infectious diseases, such as three novel pathogenic human coronaviruses in the past two decades: severe acute respiratory syndrome coronavirus 2 (SARS-CoV-2), middle eastern respiratory syndrome coronavirus (MERS-CoV) and SARS-CoV, have caused high mortality and unprecedented social and economic consequences^1–4^. While vaccines are effective in blocking infectious diseases, antibody therapy is an alternative treatment strategy for preventing newly emerging viruses. During the outbreaks of SARS-CoV, MERS-CoV and SARS-CoV-2, convalescent plasma from these patients containing neutralizing mAbs was a safe and effective treatment option to reduce mortality in severe cases^5,6^. However, convalescent plasma are limited and polyclonal non-neutralizing Abs in the plasma may cause undesired side effects^7^. The neutralizing mAbs therapeutics are effective replacements of convalescent plasma therapy. A rapid and efficient neutralizing Abs screening method against infectious diseases is in great needs.

The outbreak of COVID-19 was caused by SARS-CoV-2. Its viral spike(S) containing the receptor-binding domain (RBD) is responsible for binding to the receptor angiotensin-converting enzyme-2(ACE2) receptor on the host cells^8,9^. To date, several teams have promptly developed some potent neutralizing mAbs to SARS-CoV-2^10^–^22^. In these studies, different methodologies were employed for screening of neutralizing Abs. Some studies utilized SARS-CoV-2 S-or RBD-labeled memory B cells from convalescent patients with SASR-CoV-2 infection and directly amplified Ab genes by RT-PCR and nested PCR at a single-cell level^10,18^. In other studies, plasma cells were activated and expanded with stimulators and cytokines in vitro for selecting neutralizing Abs^14,19^. Humanized mice were also used to generate full human monoclonal antibodies against S protein^11,22^. Furthermore, using single-cell sequencing technology in combination with the enrichment of antigen-specific B, some researchers quickly detected thousands of antigen specific mAbs sequences^17,21^. Although these studies showed that neutralizing antibodies against SASR-CoV-2 could be obtained from convalescence patients, few methodologies were reported in detail about how to generate neutralizing Abs of interest rapidly and efficiently.

Here, we describe a strategically optimized system for fast screening of neutralizing mAbs to achieve efficient and reliable yield of desired neutralizing Abs in as short as 6 days. A total of 198 Abs against RBD of SARS-CoV-2 were obtained with this method, and 50% of them were potential candidate neutralizing Abs in a preliminary screening of pseudovirus system. Furthermore, 20 of these neutralizing mAbs are confirmed with high potency, and 2 mAbs reached IC_50_ in nanogram range. Therefore, the screening system can generate a large number of neutralizing mAbs for the discovery of potent antibody drugs, providing vital information to understand the characteristics of new viruses, and, collectively, to develop preventative and therapeutic strategies for newly emerging infectious diseases in the future.

## Results

### Establishment of a rapid and efficient neutralizing Abs screening system

Previously we had obtained specific mAbs from PBMC of vaccinated volunteers via microwell array chips within only one week ^23^. Here, we established the optimized screening system based on the memory B cells from the PBMC to obtain neutralizing mAbs rapidly and efficiently (Figure 1). At first, we collected the blood samples and isolated the PBMC from a panel of Chinese convalescent patients infected with SARS-CoV-2 in February. RBD specific memory B cells (mB cells) in a pooled PBMC from ^5–7^ blood samples were detected by labeling with RBD. RBD-specific mB cells were sorted into 96-well PCR plates in a single-cell manner. Each single-cell Ab cDNA was amplified, and the immunoglobulin heavy (IGH) and light chains (IGK and IGL) of the variable region were obtained by RT-PCR and nested PCR with the optimized primers at Day 1 (Table S4. Primers List of BCR PT-PCR). Recombinant sites were introduced by the nested primers during the 2^nd^ PCR. Next, linear antibody gene expression cassettes were assembled by overlapping PCR, which contained the essential elements for Ab gene transcription, including the CMV promoter, the antibody variable region, the constant region and the poly(A) tail. Then HEK293T cells were transiently transfected with these linear antibody gene expression cassettes, for the expression of recombinant antibodies at Day 2. Culture supernatants of the transfected cells were evaluated for the S and RBD specific binding activity by enzyme-linked immunosorbent assays (ELISA) at Day 4, and their pseudovirus neutralizing capacity was tested in HEK293T/hACE2 cells at Day 6. The screening system allowed us to harvest a large number of potential neutralizing Abs against SARS-CoV-2 within only 6 days. Furthermore, recombinant antibody proteins of interest were expressed and purified for subsequent functional analysis, including antibody specific binding ability, viral neutralization and antigen-binding affinity, all of which were completed with an additional 9 days.

**Figure 1.**
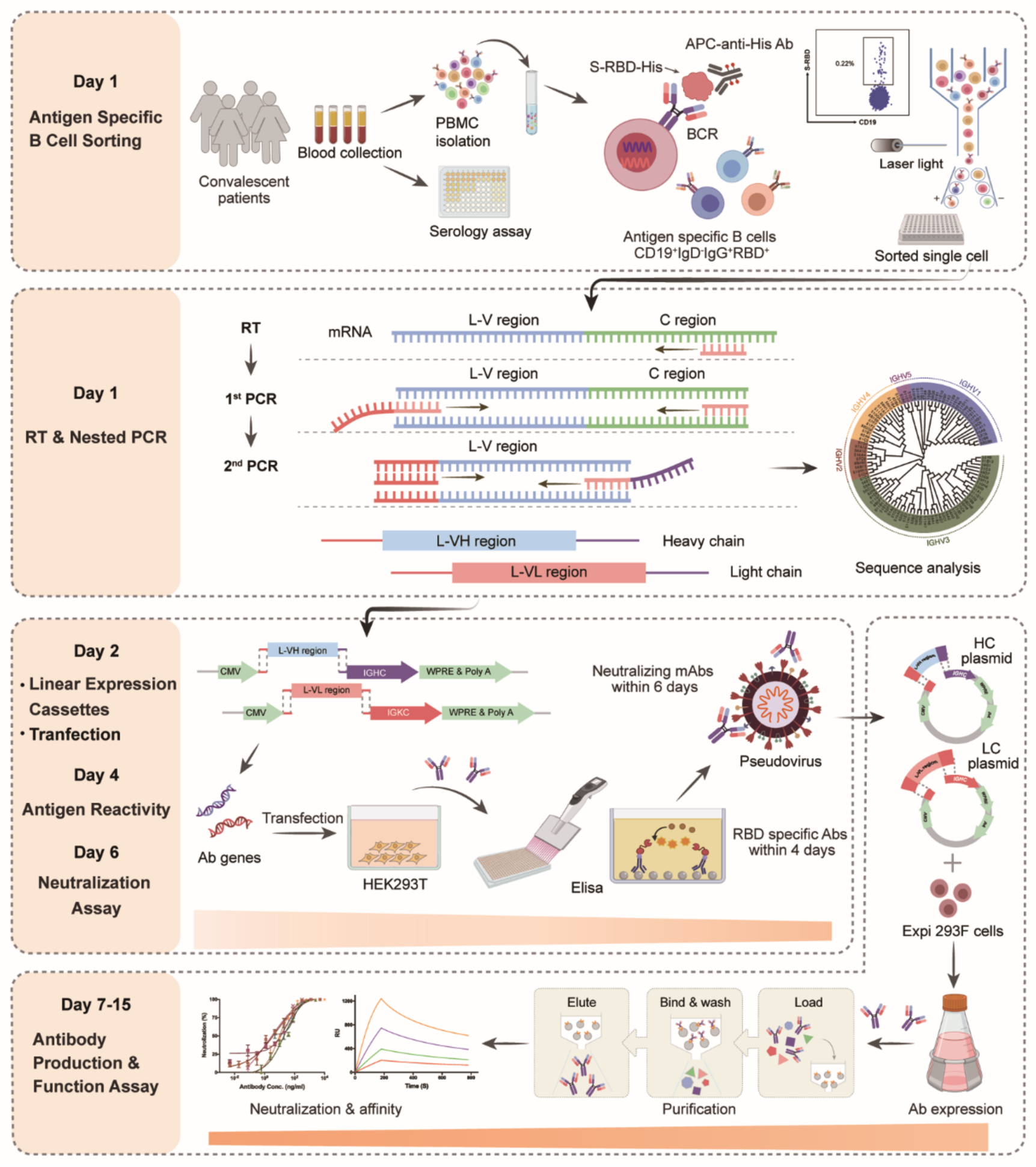
Schematic model depicting a rapid and efficient screening system of neutralizing Abs. Rapid neutralizing antibody screening workflows and timelines are shown, representing the multiple workflows conducted in parallel. PBMC were isolated from collected convalescent patients’ blood, and the RBD-specific memory B cells in the PBMCs were sorted as single-cell via flow-cytometric sorter (day 1). Then, the IgG heavy and light chains of monoclonal antibody genes were amplified by RT-PCR on the same day. 2^nd^ PCR products were cloned into linear expression cassettes on the second day. Antibodies were expressed by transient transfection with equal amounts of paired heavy and light chain linear expression cassettes in HEK293T cells and culture for two days. The cell supernatants in HEK293T cells were detected for the specificity of antibodies by ELISA in 384-well plates on the fourth day. The neutralizing activity of antibodies was detected with pseudovirus bearing SARS-CoV-2 S in 96-well plates on the sixth day. The potential neutralization antibody expression plasmids were transfected into Exi293F cells for large-scale production of Ab proteins. The cell supernatants in Exi293F cells were collected, and antibody proteins were purified by protein G. They were further measured for the binding ability and neutralizing activity via ELISA and competitive ELISA *in vitro*. Additionally, virus neutralization assay was performed. Created with Biorender.com.

Compared with conventional methods for screening neutralizing Abs^10,13,14,17,21^, a few strategically optimized details of our screening system were described as following (Figure S1. The optimization of the screening platform). First, we collected the blood samples from convalescent patients with COVID-19. These patients were the earliest confirmed cases of SARS-CoV-2 infection in Chongqing City, over half of the patients who had a history of close contact with the earliest infectors. Antigen specific B cells from memory B cells experienced affinity maturation and somatic hypermutation^24,25^. Thus, RBD specific memory B cells in these patients were potentially useful candidates to screen out the potent neutralizing Abs. Second, we sorted the CD19^+^IgD^−^IgG^+^ memory B cells in the pooled PBMC samples from multi-patients to achieve a higher probability in antibody diversity. We then utilized RBD of SARS-CoV-2 as the bait to label the specific memory B cells. Removing dead cells was essential for sorting of the relatively rare RBD specific mB cells, which only occupied less than 1% of CD19^+^IgD^−^IgG^+^ memory B cells (Figure S2. The influence of dead cells on the sorting of RBD-specific memory B cells). Additional improvements were applied to the single-B cell receptor (BCR) cloning^26,27^ and expression (Figure S3. Schematic diagram of BCR RT-PCR and linear expression cassettes construction). We chose the initial 20 nucleotides located at 5’ end of the signal peptide in Ab genes as the forward primers in the 1st PCR step, and the adaptor primer in the 2^nd^ PCR step. This was beneficial for reducing potential loss of BCR clones caused by SNP at the primer binding sites. Besides, the 2^nd^ PCR products containing the adaptor primer could be used for the simultaneous construction of linear gene expression cassettes and plasmids without the extra-modification. with Ab J-region primers in the 2^nd^ PCR, we are able to improve the recombination efficiency of linear cassettes approaching 100%. Our strategy of linear gene cassettes skipped the process of plasmid construction, which could reduce time-consuming procedures and labor, and be more suitable for a large scale of antibody screening^28^ (Table S6. The annotation of linear antibody expression cassettes). Taken together, we optimized a methodology for the rapid and efficient identification of neutralizing Ab candidates (Figure S3A, Table S3–S6).

### Detection and isolation of RBD specific memory B cells

To apply our established screening system of neutralizing Abs, we collected the plasma and PBMC from 39 convalescent patients with COVID-19 admitted to Chongqing Medical University affiliated Yongchuan Hospital (Table S1. Patient Information). These convalescent plasma had been preliminarily screened for positive virus-specific binding and neutralization capacity, using a magnetic chemiluminescence enzyme immunoassay (MCLIA) and a pseudovirus-based assay^29^. Using ELISA assay, we confirmed that those Abs targeting Spike or recombinant RBD of SARS-CoV-2, SARS-CoV, and MERS-CoV in the plasma with 10-fold dilution. Among these convalescent patient samples, 36 plasma showed high reactivity to SARS-CoV-2 S or RBD proteins, while the other three patients had weak reactivity to these antigens (Figure 2A). Almost all samples had cross-reactivity to the S1 protein of SARS-CoV and MERS-CoV with 10-or 100-fold dilution, while the healthy donor’s plasma react to none of these three coronaviruses (Figure 2A). With these findings, we felt confident that all samples could be used for the specific mAb isolation.

**Figure 2.**
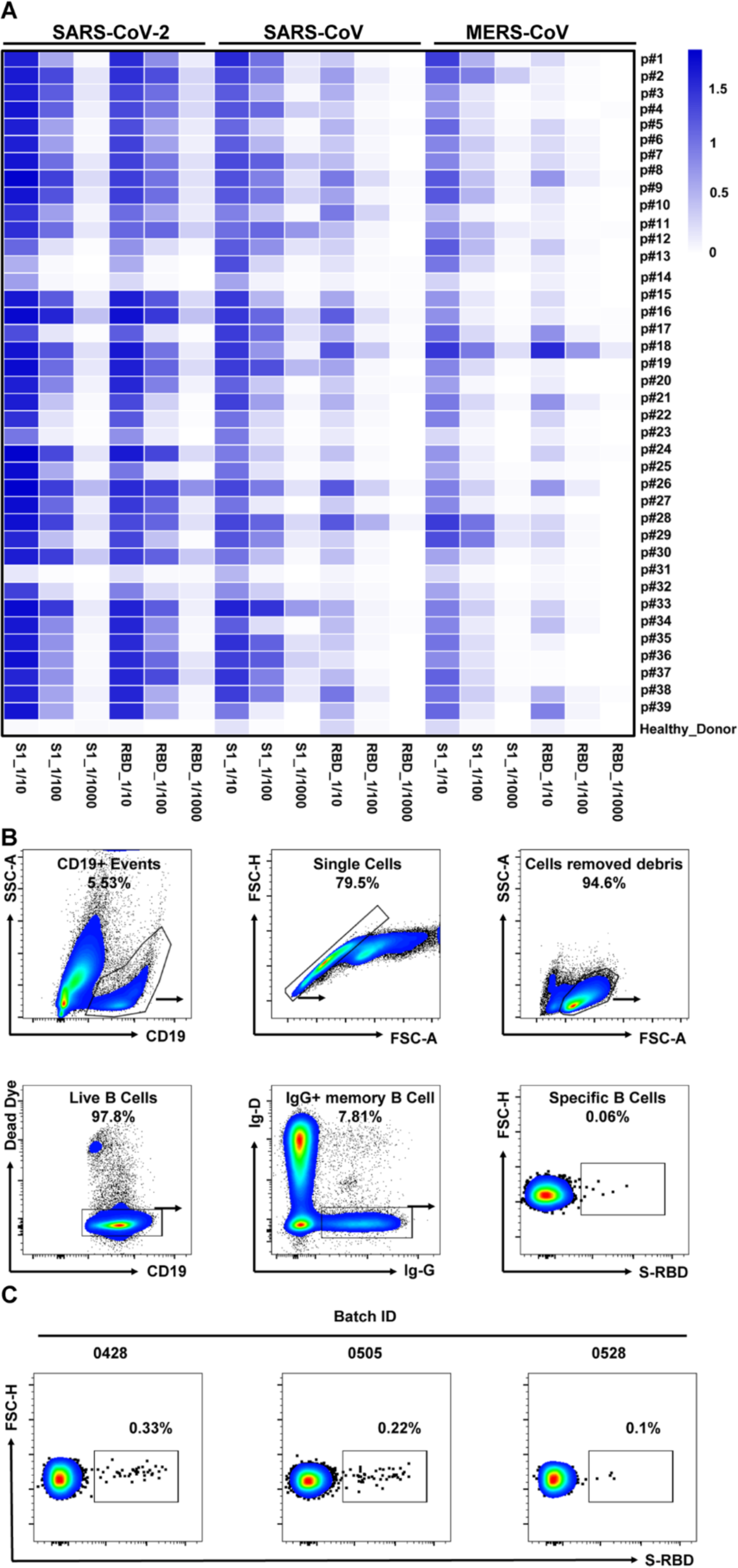
Isolation of RBD-specific memory B cells using flow cytometry. **A.** The heatmap depicts the specificity of convalescent patients’ plasma against S1 and RBD from SARS-CoV-2, SARS-CoV and MERS-CoV, measured by ELISA. Serial dilutions of plasma samples were performed to test the reactivity of antibodies in plasma. The plasma of healthy donors was used as the control. Data were shown with the mean of representative experiments. **B.** Gating strategy for SARS-CoV-2 RBD-specific IgG^+^ B cells in PBMCs of the convalescent patients. Living CD19+ IgD^−^IgG^+^ cells were gated, and cells with positive SARS-CoV-2 RBD staining were selected for single-cell sorting. **C.** FACS analysis of RBD-specific memory B cells in CD19^+^IgD^−^IgG^+^ memory B cells from PBMCs of three batch convalescent patients. Plots show CD19^+^IgD^−^IgG^+^RBD+ populations using gating strategy described in **B**.

Since RBD is the key domain for the SARS-CoV-2 S protein to interact with human cell surface ACE2 receptor, the recombinant RBD was employed to detect the specific memory B cells via flow cytometry. We analyzed RBD-specific memory B cells by a gating strategy of Dead_Dye-CD19^+^IgG^+^IgD-RBD^+^cells (Figure 2B), the proportion of which was less than 1% in IgD^−^IgG^+^ memory B cells (ranging from 0.1% to 0.33%, Figure 2C). These RBD-specific mB cells were then sorted into 96-well plate one cell per well for Ab gene isolation. Immunoglobulin heavy and light chains were amplified by nested PCR from the sorted single mB cells (Figure S3B). The amplified products were cloned into linear expression cassettes to produce full-length IgG1 antibodies (Figure S3B). After three rounds of screening, a total of 497 paired heavy chains and light chains of Ab genes were obtained from the sorted RBD-specific memory B cells (Table S2. Three batches of S-RBD specific B memory cell sorting).

### Specificity and neutralization of Abs expressed by linear expression cassettes

The amplified products with heavy chains and light chains of Ab genes were separately cloned into linear expression cassettes. We then transfected HEK293T cells with these linear expression cassettes to identify the specificity of these Abs. Antibodies within the supernatant of transfected HEK293T cells were screened by ELISA for their binding capability to the recombinant S1 and RBD protein of SARS-CoV-2. In total, we identified 198 RBD specific antibody genes from the 497 pair Ab genes (Figure 3A). To assess the neutralization ability of these specific antibodies, we used pseudovirus bearing SARS-CoV-2 S protein to infect 293T/hACE2 cells. Interestingly, 96 out of 198 antibodies (48.5%) showed the potential ability to block pseudovirus with an inhibitory rate of over 75% (Figure 3A), suggesting that RBD region was an ideal candidate to screen neutralizing Abs for blocking SARS-CoV-2. These results demonstrated that our screening system can rapidly and efficiently screen neutralizing Abs using patients’ specific memory B cells.

**Figure 3.**
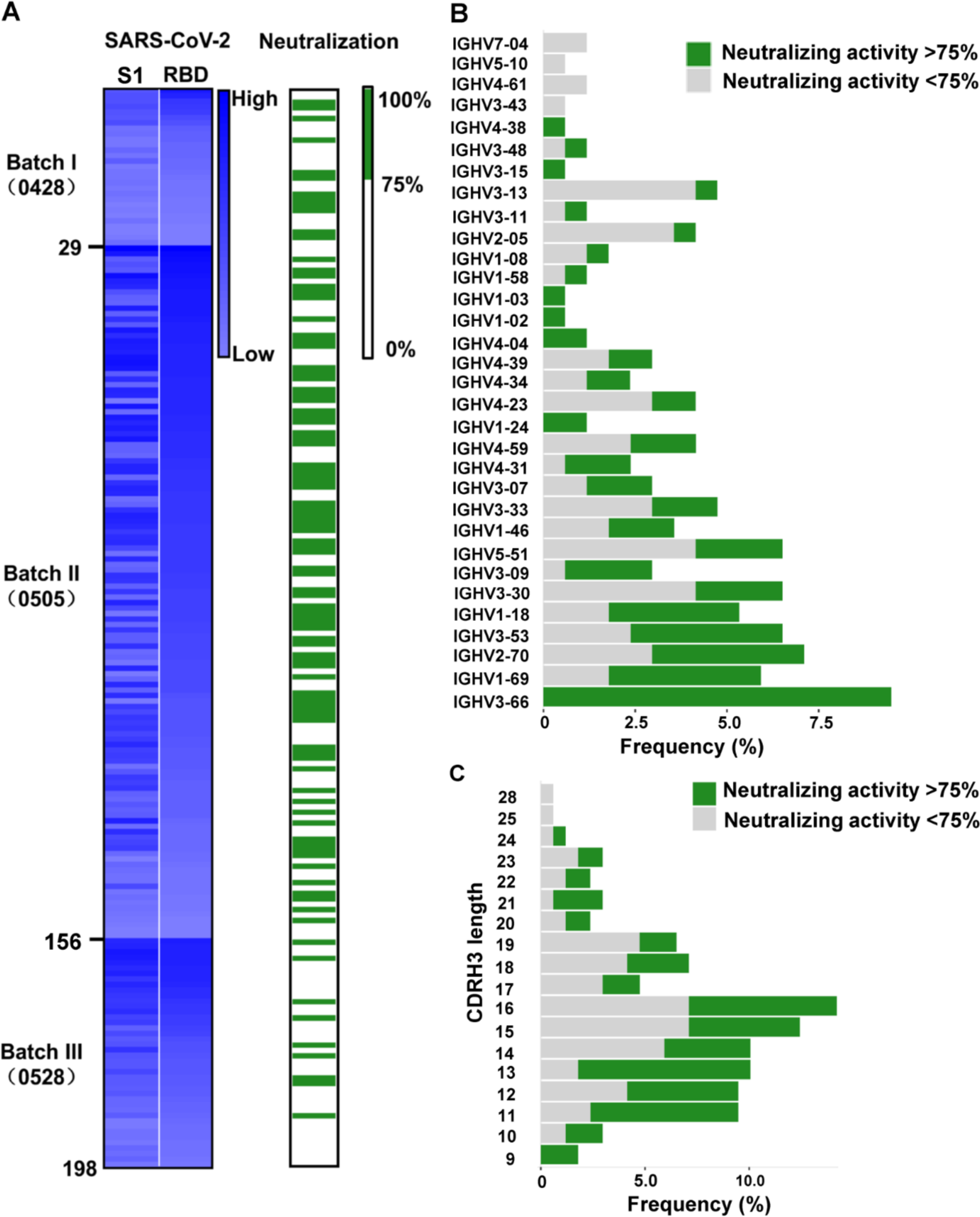
Identification of RBD specific monoclonal antibodies from convalescent COVID-19 patients. **A.** Screening of specific Abs against SARS-CoV-2 S1 and RBD. The heatmap reveals that the binding ability of 198 Ab supernatants produced by HEK239T cells transfected with linear Ab gene expression cassette. The mAbs rank as the screening sequence, and binding activity of mAbs against SARS-CoV-2 S1 and RBD were tested by ELISA. The brightness of blue represents the binding strength, which reflected the OD405 nm value tested by ELISA. The neutralizing activity of mAbs was discriminated according to the neutralizing value. Antibody-mediated blocking of luciferase-encoding SARS-CoV-2 typed pseudovirus transfected into hACE2/ HEK293T cells were measured by values of relative light units (RUL). The Green columns indicate potential neutralization (neutralizing activity >75%), while white indicate partial or not neutralization (neutralizing activity <75%). **B.** Frequencies of variable region of heavy chain (VH) gene clusters for potential neutralizing and non-neutralizing antibodies. Clonal sequences groups were collapsed and treated as one sample for calculation of the frequencies. **C.** Frequency of various the heavy chain complementarity determining region 3 (CDRH3) length of in potential neutralizing and non-neutralizing antibodies.

### Sequence analysis for the diversity of RBD-specific Abs

We then successfully sequenced 169 RBD-specific Abs. Among them, 158 (93.5%) Abs had unique patterns of distribution with various gene clusters (Figure S4. Usage and pairing of heavy and light chain for all specific antibodies). We also analyzed the distribution of heavy chain and light chain gene clusters according to neutralizing capability of mAbs tested by pseudovirus assays, as shown in Figure 3B and Figure S4A. Abs with neutralizing rate over 75% to block pseudovirus were termed as potential neutralizing Abs (pote-nAbs). The almost sequenced Abs were transcribed from the IGHV1-IGHV5 for the heavy chain and IGKV1-IGKV3 and IGLV1-IGLV3 for light chain (Figure S5). We found that close to 50% of the pote-nAbs were specifically transcribed from IGHV3 for the heavy chain, and from IGKV1 for the light chain (Figure S5. Phylogenetic analysis of VH (up) and VL (down) genes for RBD-binding antibodies). Interestingly, we found that all mAbs encoded by IGHV3-66 were pote-nAbs (Figure 3B), and the IGHV3-66 family paired with IGKV1-33, IGKV1-9 and IGLV1-40 (Figure S4C). Additionally, a large number of mAbs encoded by IGKV1-39 were pote-nAbs (Figure S4A). Of note, we found that IGKV1-39 gene cluster also paired with a bundle of heavy chains to express RBD specific mAbs (Figure S4C), which was consistent with previous reports ^11,19^.

The heavy chain complementarity determining region 3 (CDRH3) is the most variable region of an antibody in amino acid compositions and lengths. The average length of CDRH3 in the naive human repertoire is round 15 amino acids with a normal distribution^30^. We observed that the CDRH3 lengths of the specific mAbs were mainly distributed between 11-19 amino acids, while the overall CDRH3 lengths matched the skew distribution (Figure 3C). Most of the potent-Abs contained ^11–16^ amino acids (Figure 3C). The mean CDRH3 length of isolated SARS-CoV-2 S-RBD-specific mB cells differs substantially from that of other viral infections, such as HIV and influenza virus^31,32^. In terms of the CDR3 light chain (CDRL3) lengths, a range of 6 to 13 amino acids were observed, with similar skew distribution (Figure S4B).

### Potent neutralizing ability and antigen affinity of mAbs

The variable regions of potential neutralizing Abs were cloned into antibody expression vectors to construct Ab plasmids. We successfully harvested 73 purified mAbs, from a total of 96 potential neutralizing mAbs that were produced by transfecting Expi293F cells. When we tested the specificity of purified mAbs by ELISA, we found 65 Abs that formed tight interaction with SARS-CoV-2 S1 and SARS-CoV-2 RBD (Figure 4A). Next, these mAbs were assessed with RBD-ACE2 interaction blocking assay to confirm their neutralizing ability *in vitro*. We found that 71% of them could block the interaction between ACE2 and RBD (Figure 4B).

**Figure 4.**
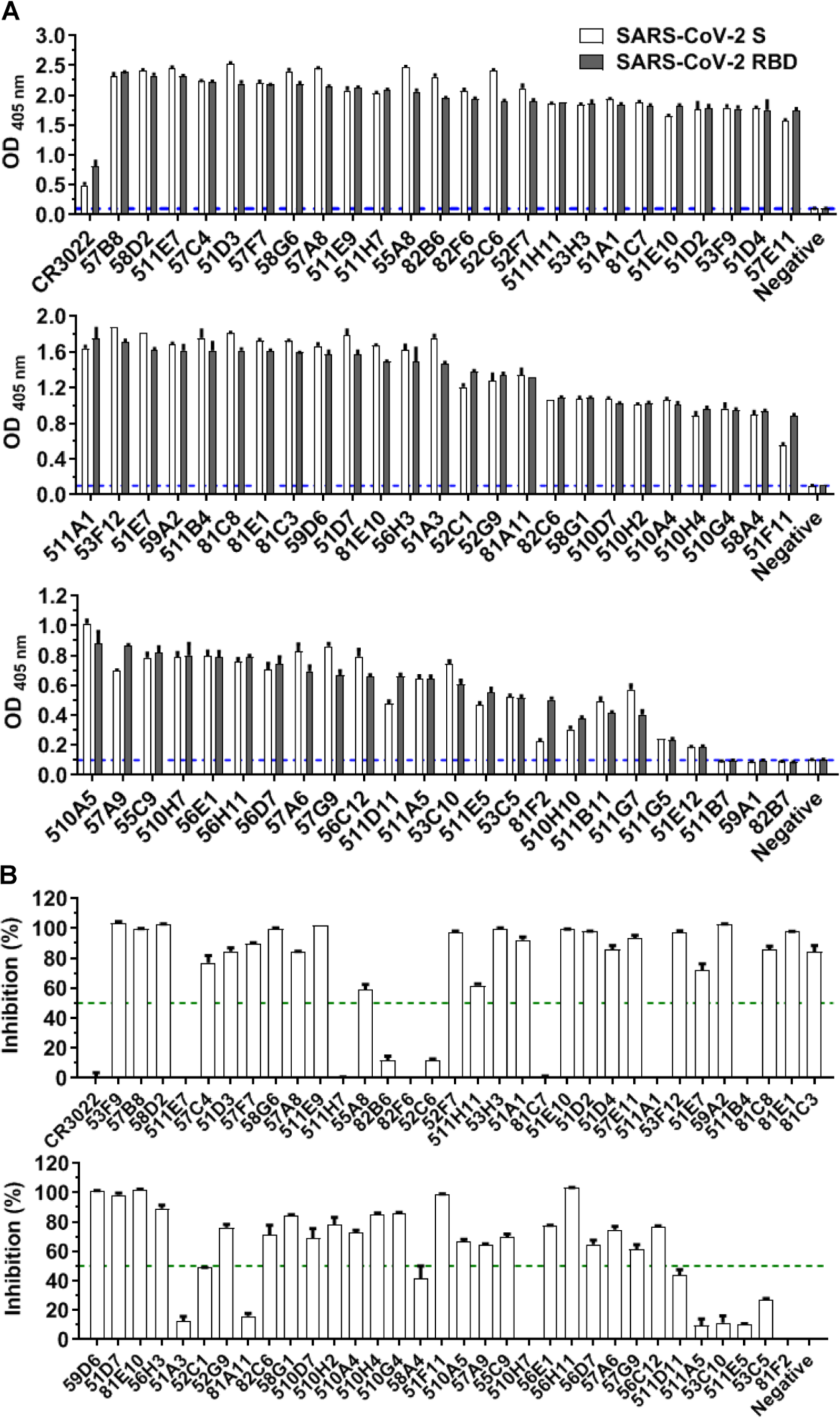
The binding activity and inhibition of ACE2-RBD interaction of mAbs tested by ELISA and competitive ELISA. **A.** The OD_405 nm_ value refects a binding strength of purified mAbs to 1 μg/ml SARS-CoV-2 S1 or RBD. Plates were coated with recombinant S1 or RBD protein of SARS-CoV-2, then incubated with purified mAbs. A SARS specific mAb (CR3022) was set as the positive control. The blue dashed lines indicated the OD_405nm_ value of a negative sample. **B.** The inhibitory effect of purified mAbs against the interaction between SARS-CoV-2 RBD and hACE2 was tested via competitive ELISA analysis. Blocking efficacy was determined by comparing response units with and without prior antibody incubation. The green dashed lines indicated 50% inhibition on blocking the interaction ACE2 and RBD interaction.

Forty-eight purified mAbs were evaluated for their neutralizing potency using the authentic SARS-CoV-2 cytopathic effect (CPE) inhibition assay, and the results were listed according to the order of inhibitory potency (Figure 5A). We successfully obtained a total of 20 antibodies that were able to completely block the authentic SARS-CoV-2 infection with the concentrations of 1 μg/ml. The top level 2 mAbs on the list were termed as the most potent neutralizers (completely inhibition < 0.14 μg/ml), and another 18 mAbs as the moderate neutralizers (0.29-1.17 μg/ml). The IC_50_ of the most potent neutralizers (58G6, 510A5) were determined by RT-qPCR method using authentic SARS-CoV-2 virus infection. We found that the IC_50_ values of 58G6 and 510A5 were 9.98 ng/ml and 11.13 ng/ml, respectively (Figure 5B). Therefore, we tested the binding affinity of 58G6 and 510A5 to SARS-CoV-2 S-RBD via the surface plasmon resonance (SPR) assay. The measured equilibrium constant (Kd) of 58G6 with SARS-CoV-2 S-RBD was 0.385 nM and that of 510A5 was 7.8 nM, respectively (Figure 5C). In our study, 58G6 and 510A5 are the best mAsb with potent neutralization and high affinity against SARS-CoV-2.

**Figure 5.**
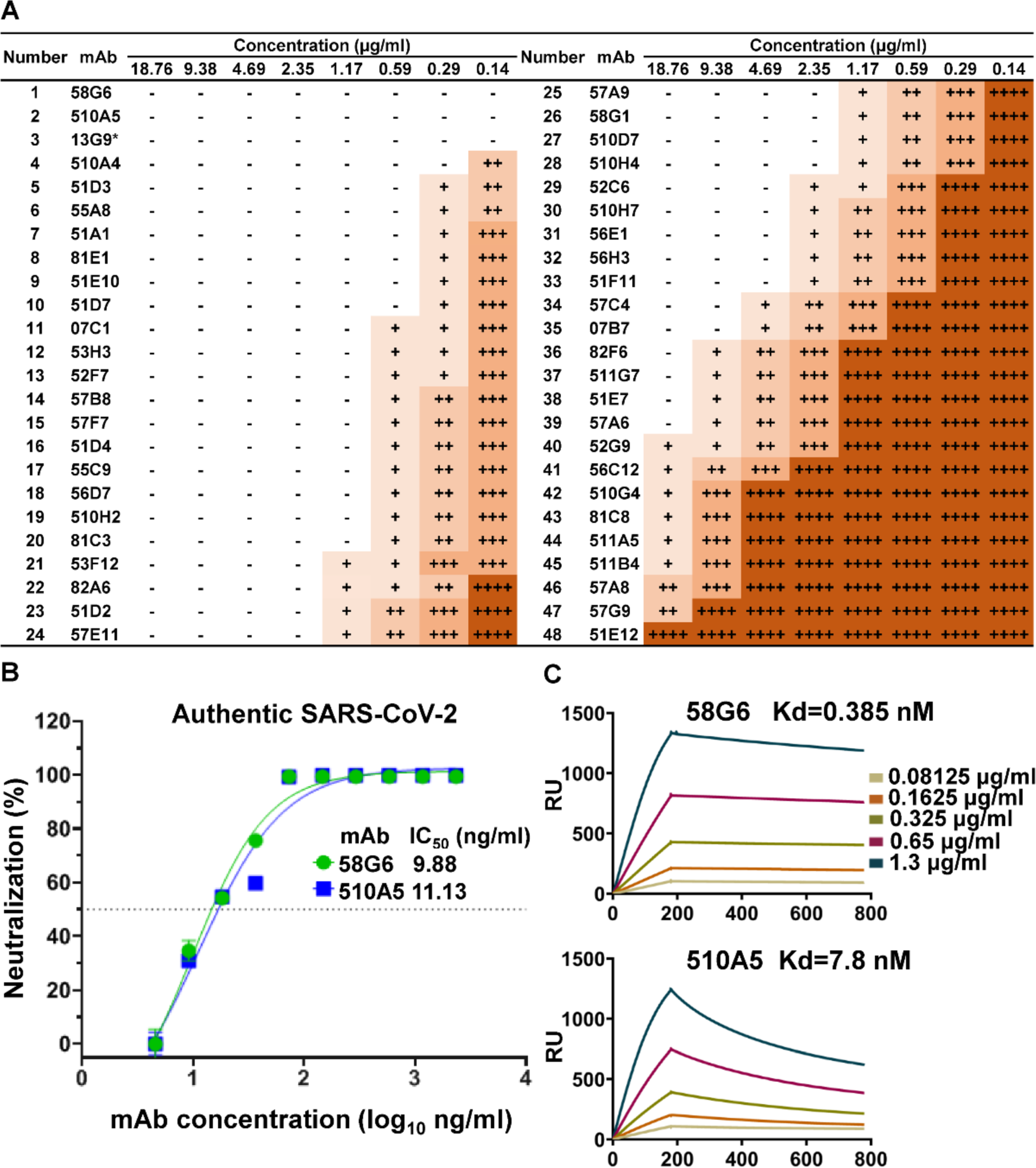
Functional characteristics of neutralizing Abs against SARS-CoV-2. **A.** Neutralization activity of mAbs against authentic SARS-CoV-2 virus (nCoV-SH01) were analyzed by Cytopathic effects (CPE) test. Serial dilutions of mAbs were tested in parallel against authentic SARS-CoV-2, ranging from 18.76 μg/ml to 0.14 μg/ml. CPE results was summarized in (A) where “++++” indicates 100% cytopathy, “+++” indicates 50-75%, “++” indicates 25-50%, “+” indicates <25% and “-” indicates no cytopathy. 13G9 was marked “*”, which was obtained by the method previously described^23^. **B.** The neutralization activity of 58G6 and 510A5 against the authentic SARS-CoV-2 virus was determined in Vero-E6 cells by RT-qPCR. Dashed lines indicated a 50% reduction in viral infectivity. Data were shown as mean ± SD of representative experiments. **C.** Binding kinetics of isolated mAbs with SARS-CoV-2 RBD were measured by Surface Plasmon Resonance (SPR). The purified antibody was captured onto the CM5 sensor chip, followed by the injection of soluble SARS-CoV-2 RBD at five different concentrations. The experimental data of 58G6 and 510A5 were shown in the top and bottom figures in **C** respectively. The results presented are representatives of two independent experiments.

## Discussion

Neutralizing antibodies were considered as an ideal medicine for prophylaxis and treatment of infectious diseases^33^. At present, several potent neutralizing Abs to SARS-CoV-2 have been promptly developed and being tested in clinical trials (clinicaltrials.gov NCT04497987, NCT04426695 and NCT04425629) for treating COVID-19 patients ^12–15,17,19–21,34,35^. These reports showed that even though neutralizing antibodies against SASR-CoV-2 could be obtained from convalescent patients, the success rate to discover potent neutralizing antibodies with therapeutic value remains unideal. In this study, we described a strategically optimized screening method to discover potent mAbs from a large number of potential neutralizing Abs.

In general, here is how researchers obtain neutralizing antibodies. Blood samples were collected from convalescent patients with SASR-CoV-2 infection, and PBMC were separated in order to isolate SASR-CoV-2 specific B cells. Then the paired heavy and light chain sequences of Ab genes were obtained either in a single-cell PCR manner^10,12^,^15,18^, or directly by high-throughput single-cell sequencing^17,21^. In our study, we obtained the paired Ab genes at the single-cell level, and optimized steps of a screening workflow for neutralizing Abs, we were able to obtain the most potent neutralizing Abs with high speed and efficiency.

Firstly, we optimized the specificity of antibody isolation to increase target neutralizing Abs probability. Because the key sites of RBD have been clearly demonstrated to be essential for ACE2 binding during SARS-CoV-2 entry^8,36^, we chose to sort RBD-specific memory B cells for the isolation of heavy and light chains. It has been evidenced that RBD-specific neutralizing Abs can inhibit SARS-CoV-2 entry to host cells^10^, we have observed similar findings with psudovirus infection (Figure 3). Recent reports have demonstrated that when SARS-CoV-2 S was used as bait to label antigen-specific mB cells, it could result in a large of undesired antibodies against over-broad antigenic sites belonging to non-RBD regions^12,17^. When we used RBD as bait to isolate antibodies from mB cells, we found the obtained mAbs were more effective in blocking authentic viruses (Figure 5), which might largely due to the fact that RBD was enriched with the ACE2 binding epitopes.

Secondly, we optimized multiple steps to improve the efficiency of nAb screening. This has been achieved by smarter primer design, application of linear expression cassettes and preliminary neutralization assay to exclude non-neutralizing Abs, and these will be discussed in detail. Initially, we designed primers targeting the initial 20 nucleotides at the 5’ end of the signal peptide of Ab genes as the forward primers in the 1st PCR step. This can reduce the loss of BCR clones caused by SNP at the primer binding sites. Also, we added an adaptor primer in the 2^nd^ PCR step. Such adaptor with the same sequences as downstream recombination sites was convenient for the next PCR and recombinant plasmids construction, which could be suitable for high-throughput screening of specific Abs. Next, we successfully constructed linear expression cassettes with heavy chains or light chains, to rapidly identify the specificity of Abs. Construction of the linear expression cassettes was much easier than plasmids, which could drastically reduce the workload and time. Moreover, the linear Ab gene expression cassettes expressed in the cell supernatants of HEK293T cells were evaluated for neutralization activity on the sixth day. Neutralizing Abs account for approximately 50% of RBD-specific Abs, as shown in Figure 3A. By detecting the ability of purified mAbs to block the interaction of RBD with ACE2, we could filter out only those Abs with neutralizing activities, to be applied in the subsequent steps. Together, these optimizations allowed us to finish one round of the screening for neutralizing Abs in, as short as, 15 days. The details of this optimized steps of our established methodology are shown (Figure S3, Table S03~S05). Thirdly, we used both authentic SARS-CoV-2 cytopathic effect (CPE) inhibition assay and quantitative analysis by RT-qPCR to ensure the accuracy of our findings, which lead to the successful identification of 20 potent neutralizing antibodies that can completely block authentic virus infection, at concentrations 1.17 μg/ml. And the top two antibodies (58G6 and 510A5) generated IC_50_ values at around 10 ng/ml, which were, as far as we know, two of the most potent neutralizing mAbs discovered to date.

Last but not least, we reduced duplicating clones by sample selection and increased the efficiency of BCR cloning by adjusting gating strategy. It has been reported that substantial mAbs clones to SARS-CoV-2 expanding in the individual patient sample is relatively common^13,18^. In our study, almost all RBD-specific mAb clones were different from one another, while the proportion of clonal expansion was only 6.5% in all sequences, as compared to approximately 20%^13,18^. It could result from our pooled PBMC for sorting RBD-specific memory B from 5–7 convalescent COVID-19 patients. Therefore, it is beneficial to improve sample selection by methods that can best yield diverse mAbs of interest. Furthermore, RBD-specific memory B cells were mainly sorted after removing dead cells, this process increased the efficiency of BCR cloning.

Such optimized screening system allowed us to efficiently generate a panel of neutralizing Abs with relatively great potency. When we analyzed the distribution of gene clusters of B cell receptor (BCR) repertoire of abundant potential neutralizing and non-neutralizing antibody sequences, a few interesting observations were found. Our results revealed that potential neutralizing Abs tended to be distributed in several gene clusters, such as VH3-66 and VH3-53 allele, etc., among which, the VH3-66 has exclusively produced neutralizing Abs. This result may be helpful in analyzing the preference distribution of neutralizing Abs in the future. Meanwhile, CDRH3 length is reported as a key factor to value the diversity of RBD specific Abs, due to the changeable amino acid composition. We found that the CDRH3 length of potential neutralizing Abs showed a skewed distribution, with an inclined length of ^11–16^ amino acids. It suggested that the SARS-CoV-2 antibodies were likely derived from memory B cells during the primary response to SARS-CoV-2 infection but not a recall response to SARS-CoV or MERS, even though our collected blood specimens were cross-reactive with both SARS-CoV and MERS S protein32. One additional improvement that can be integrated into our screening system is single-cell sequencing. The development of proper algorithms for neutralization evaluation with incorporation of heavy chain variable region preferences, for example, IGHV3-66, could help to precisely predict neutralizing antibody from thousands of antigen specific mAbs repertoire^17,21^. This might further provide desired candidates of neutralizing Abs with potential therapeutic value, with better time-efficiency and economical preferences.

In conclusion, we have successfully established a strategically optimized screening system of neutralizing antibodies that can generate ideal numbers of neutralizing Abs in a total period of 15 days. This methodology can open the way for the potential on-time therapeutic applications towards various emerging pathogens in the future.

## Materials and Methods

### Isolation of single RBD-specific memory B cells by FACS

PBMCs from the convalescent patients were thawed and rested overnight. The mixed samples staining as following. 2 μg/ml RBD-his in 200 μl PBS (added with 2% FBS) was mixed with the specific antibody cocktail required for staining B cell. Then these PBMCs was incubated with mixed antibodies cocktail at 4 °C for 30 min (the antibodies cocktail including FITC-anti-human CD19 antibody (Biolegend, clone: SJ25C1), BV421-anti-human IgD antibody (Biolegend, clone: IA6-2), PerCP-Cy5.5-anti-human IgG antibody (Biolegend, clone: M1310G05), APC-anti-his tag antibody (Biolegend, clone: J095G46)). Dead dye (LIVE/DEAD™ Fixable Near-IR Dead Cell Stain Kit, Thermo Fisher) was added at 4 °C for 20 min. After washing the cells, the FACS analysis were performed by BD FACSAriaIII (BD Biosciences) with FSC-A versus SSC-A identifying cell population, FSC-A versus FSC-H excluding doublets. Then FSC-H versus Dead Dye was gated to remove dead cells. RBD-specific single memory B cells were gated by CD19^+^IgD^−^IgG^+^His^+^, and single-cell sorted into 96-well PCR plates (free of DNase and RNase, Bio-Rad). The Plates were stored at −80 °C until BCR Cloning. Data analysis was performed utilizing the FlowJo software (FlowJo, LLC).

### Amplification of single-cell BCR variable region

Our primers for PCR were designed from leader sequences and J region sequence of immunoglobulin (Ig) annotated by the IMGT reference directory (http://www.imgt.org/vquest/refseqh.html). An adaptor sequence was added to the 5′ end of the leader primers for the 2^nd^ PCR. 31 leader primers (AP_G_leader Mix) was designed for the heavy chain of Ig, and 19 leader primers (AP_K_leader Mix) was used in the amplification of the kappa chain of Ig, and 21 leader primers (AP_L_leader Mix) for the lambda chain of Ig were designed. For the initial step of RT-PCR, 5 μl of the RT_Mix_A was added into each well of 96 well plate containing a single B cell. Then the mixture was incubated at 65°C for 5 min and put on ice immediately for 3 min. 5 μl RT_Mix_B was added into each well of the plate with reaction program: 45 °C for 45 min, 70 °C for 15 min. 1 μl of RT product was moved to the well of a new 96 well plate containing 9 μl 1st_PCR_Mix_Gamma /Kappa/Lamda, respectively. The PCR program for 1st PCR: 95°C for 3 min, 30 cycles of 95°C for 10 sec, 55°C for 5 sec and 72°C for 1 min. 1 μl of the tenfold-diluted 1st PCR product was then added into each well of a new 96 well plate holding 9 μl 2^nd^ PCR_Mix_Gamma/Kappa/Lamda, respectively. The PCR program for 2^nd^ PCR: 95°C for 3 min, 35 cycles of 95°C for 10 sec, 55°C for 5 sec, and 72°C for 45 sec. The second PCR products were further cloned into the antibody linear expression cassettes or expression vectors to express full IgG1 antibodies. PCR reaction Mix are prepared as described in Table S3. All of the PCR primers are listed in Table S4 and prepared in Table S5.

### Generation of linear antibody expression cassettes and expression of Abs

2^nd^ PCR products were used to ligate with the expression cassettes directly by overlapping PCR. The products were purified with ethanol precipitation method. Briefly, 120 μl of absolute ethanol and 6 μl of 3 M sodium acetate were mixed with 60 μl of the Overlap PCR product. Then the reagents were incubated at −80 °C for 30 minutes. After centrifuging at 10,000 rpm for 20 minutes, the supernatant was discarded the and the pellet adhered on the tube were rinsed with 200 μl 70% ethanol and absolute ethanol and evaporated the ethanol at 56°C for 10 min. 40 μl sterile water was added to dissolve the DNA pellet. After measuring the nucleic acid concentration, purified overlapping PCR products of paired heavy and light chain expression cassettes were co-transfected in HEK293T cells. The binding ability of transfected culture supernatants to SARS-CoV-2 S-RBD was tested by ELISA after 48 hours.

### Recombinant antibody production and purification

For the construction of antibody expression Vectors, VH and VL 2^nd^ PCR products were inserted separately into the linearized plasmids (pcDNA3.4) that encode constant regions of the heavy chains and light chains via a homologous recombination kit (Catalog No. C112, Vazyme). A pair of plasmids separately expressing heavy and light chain of antibodies were transiently co-transfected into Expi293™ cells (Catalog No. A14528, ThermoFisher) with ExpiFectamine™ 293 Reagent. Then the cells were cultured in shaker incubator at 120 rpm and 8% CO2 at 37 °C. After 7 days, the supernatants with the secretion of antibodies were collected and captured by protein G Sepharose (GE Healthcare). The bound antibodies on the Sepharose were eluted and dialyzed into phosphate-buffered saline (PBS). The purified antibodies were used in following binding and neutralization analyses.

### ELISA binding assay and competitive ELISA

2 μg/ml the recombinant S or RBD proteins derived from SARS-CoV-2, SARS-CoV, or MERS-CoV (Sino Biological, Beijing) were coated on 384-well plates (Corning) at 4°C overnight. Plates were blocked with blocking buffer (PBS containing 5% FBS and 2% BSA) at 37°C for 1 hour. Serially diluted convalescents’ plasma or mAbs were added into the plates and incubated at 37°C for 30 min. Plates were washed with phosphate-buffered saline, 0.05% Tween-20 (PBST) and ALP-conjugated goat anti-human IgG (H+L) antibody (Thermo Fisher) was added into each well and incubated at 37°C for 1 hour. Lastly, the PNPP substrate was added, and absorbance was measured at 405 nm by a microplate reader (Thermo Fisher). For a competitive ELISA to test the effect of mAbs on blocking ACE2 binding RBD, 2 μg/ml the recombinant ACE2 (Sino Biological, Beijing) was added in 384-well plates and overnight at 4°C, followed by blocking with the blocking buffer and washing. 500 ng/ml RBD-mouse-Ig-Fc was pre-incubated with test specimen at 37°C for 1 hour, followed by adding into the wells coated with ACE2 and incubated at 37°C for 1 hour. Unbound antigen were removed with washes. Then ALP-conjugated anti-mouse-Ig-Fc antibody was added into the wells and incubated at 37°C for 30 min. PNPP was added and measured as above.

### Pseudovirus neutralization assay

Pseudovirus was generated as previously described^37,38^. HEK293T cells were transfected with psPAX2, pWPXL Luciferase, and pMD2.G plasmid encoding either SARS-CoV-2 S. The supernatants were harvested 48 hours later, filtered by 0.45 μm filter and centrifugated at 300 g for 10 min to collect the supernatant and then aliquoted and storied at −80°C. The purified antibodies with serial dilution were incubated with pseudovirus at 37°C for 1 hour. The mixture of viruses and specimens was then added in a hACE2 expressing cell line (hACE2-293T cell). After 48 hours culture, the luciferase activity of infected hACE2/293T cells was measured by the Bright-Luciferase Reporter Assay System (Promega). Relative luminescence units (RLU) of Luc activity was detected using ThermoFisher LUX reader. All experiments were performed at least three times and expressed as means ± standard deviations (SDs). Half-maximal inhibitory concentrations (IC50) were calculated using the four-parameter logistic regression in GraphPad Prism 8.0.

### Authentic SARS-CoV-2 virus neutralization assays

An authentic SARS-CoV-2 neutralization assay was performed in a biosafety level 3 laboratory of Fudan University. Serially diluted mAbs were incubated with authentic SARS-CoV-2 (nCoV-SH01, GenBank: MT121215.1, 100 TCID50) at 37°C for 1 hour. After incubation, the mixtures were then transferred into 96-well plates, which were seeded with Vero E6 cells. After incubation at 37°C for 48 hours, each well was examined for CPE and supernatant viral RNA by RT-qPCR. For RT-qPCR, the viral RNA was extracted from the collected supernatant using Trizol LS (Invitrogen) and used as templates for RT-qPCR analysis by Verso 1-step RT-qPCR Kit (Thermo Scientific) following the manufacturer’s instructions. PCR primers targeting SARS-CoV-2 N gene (nt608-706) were as followed (forward/reverse):5′-GGGGAACTTCTCCTGCTAGAAT-3′/5′-CAGACATTTTGCTCTCAAGCTG-3′. qRT-PCR was performed using the LightCycler 480 II PCR System (Roche) with program as followed: 50°C 15 min; 95°C 15 min; 40 cycles of 95°C 15 sec, 50°C 30 sec, 72°C 30 sec.

### Antibody binding affinity measurement by SPR

The affinity of antibody binding SARS-Cov-2-S-RBD was measured via the Biacore X100 platform. The CM5 chip (GE Healthcare) was coupled with an anti-human IgG-Fc antibody to capture 9000 response units antibodies. Gradient concentrations of SARS-Cov-2-S-RBD (Sino Biological Inc.) were diluted (2-fold dilution, from 50 nM to 0.78 nM) with HBS-EP+ Buffer (0.01 M HEPES, 0.15 M NaCl, 0.003 M EDTA and 0.05% (v/v) Surfactant P20, pH 7.4), then injected into the human IgG capturing chip. The sensor surface was regenerated with 3 M magnesium chloride at the end of each cycle. The affinity was calculated using a 1:1 binding fit model in Biacore X100 Evaluation software (Version:2.0.2).

### Sequence analysis of antigen-specific mAb sequences

IMGT/V-QUEST (http://www.imgt.org/IMGT_vquest/vquest) and IgBLAST (https://www.ncbi.nlm.nih.gov/igblast/), MIXCR (https://mixcr.readthedocs.io/en/master/) and VDJtools (https://vdjtools-doc.readthedocs.io/en/master/overlap.html) tools were used to do the VDJ analysis and annotation, germline divergence for each antibody clone. The Phylogeny tree analysis of IgG heavy and light chain variable genes was performed with MegaX (Molecular Evolutionary Genetics Analysis across computing platforms) by the Maximum Likelihood method. Abs DNA sequences were compared with each other by ClustalW (pairwise alignments) to analyze sequence similarity, and EvolView (https://www.evolgenius.info/evolview/) was used for the decoration of Phylogeny tree. R packages (ggplot2, pheatmap) were used for the bar chart, heatmap and Cicos plot.

### Ethics Statement

The project “The application of antibody tests patients infected with SARS-CoV-2” was approved by the ethics committee of ChongQing Medical University. Informed consents were obtained from all participants.

## Acknowledgments

We acknowledge the work and contribution of blood sample providers from Chongqing Medical University affiliated Yongchuan Hospital and the third affiliated Hospital of Chongqing Medical University. We also thank health donors from Chongqing Medical University. This study was supported by Chongqing Medical University fund (X4457) with the donation from Mr Yuling Feng.

## Author contributions

AJ, AH conceived and designed the study, KD, CH, LD, YN offered help on collection of convalescent patient blood samples. Most of the experiments were completed by XH,TL, CH, YW, JW, RW, FG, JH, SM, YL, FL, SS, YH, QC, LL with the assistance from TN, YX, CG, HJ, YW, WX, XC, QG, GZ, CH, WK. SL, MS, YW, XH, AJ played an import role in data analysis of neutralizing Abs sequences. SL, MS, YW, JW performed to generated figures and tables and take responsibility for the integrity and accuracy of data presention. AJ, XH wrote the manuscript and SL, TL, JW and YW helped to revise it.

## Data availability statements

All information presented in this study will be upload soon.

---

**Figure S1.**
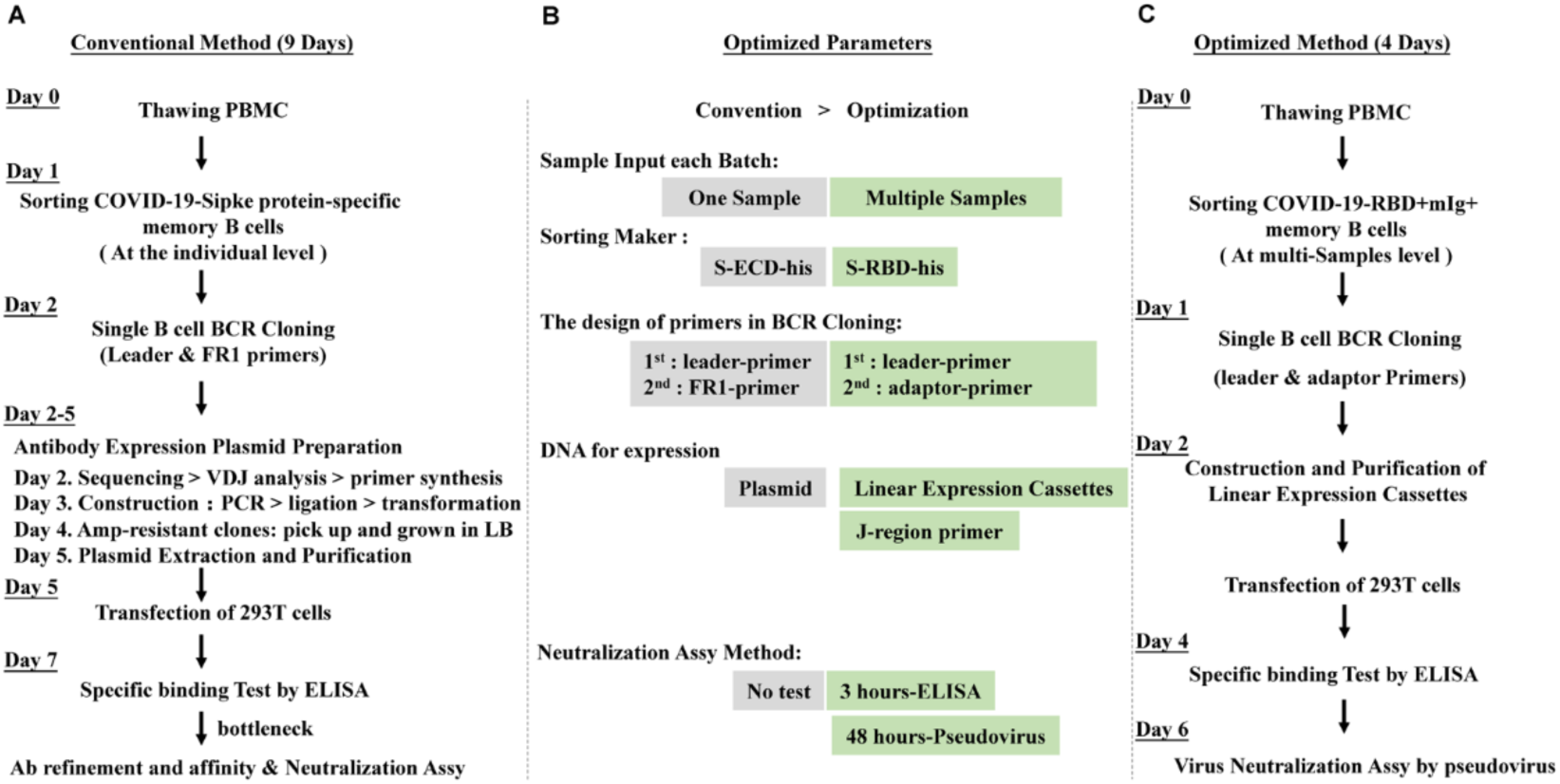
The optimization of the screening platform. **A.** The conventional screening of neutralizing antibodies. Antigen-specific B cells from PBMCs were sorted on day 1. The single-cell BCR genes were amplified by PCR on day 2. The antibody expression vectors were constructed in the next three days, including the PCR product sequencing, the primer synthesis, the ligation of genes and vectors, the DNA transformation and the plasmid extraction. The purified plasmids were transfected into HEK293T cells on day 5. After 48 hours, the cell supernatants were collected and analyzed with specific antigens by ELISA. Specific antibodies are used for following antibody expression and purification. Purified antibodies were screened as neutralizing candidates. **B.** The key parameters affecting screening efficiency. The following steps of the screening processes were carefully modified: multi-step sorting for the individual samples or the pooled samples, labeling S or S-RBD specific B cells, expressing antibodies using linear expression cassettes or plasmids, and designing preferred primers for the single-cell BCR cloning. To reduce time-consuming and workload, it is the critical step to screen neutralizing antibodies during the initial screening in the sixty days. Two methods for neutralization evaluation, competitive ELISA method in 3 hours or pseudovirus assay for 48 hours, were used side by side for confirmation of nAb neutralizing capability. **C.** The optimized strategy of neutralizing antibodies development. One day after PBMC thawing, specific B cell sorting was performed on day 1. A single BCR gene was cloned on day 2, using the 2^nd^ PCR product to construct the linear expression cassettes, which were termed as the transfection targets to be introduced directly into HEK293T cells with liposome, without constructing plasmid, to shorten the screening duration. After 48 hours, the supernatants of each transfected samples were harvested and analyzed via ELISA and pseudovirus neutralization assay for evaluating specificity and neutralization.

**Figure S2.**
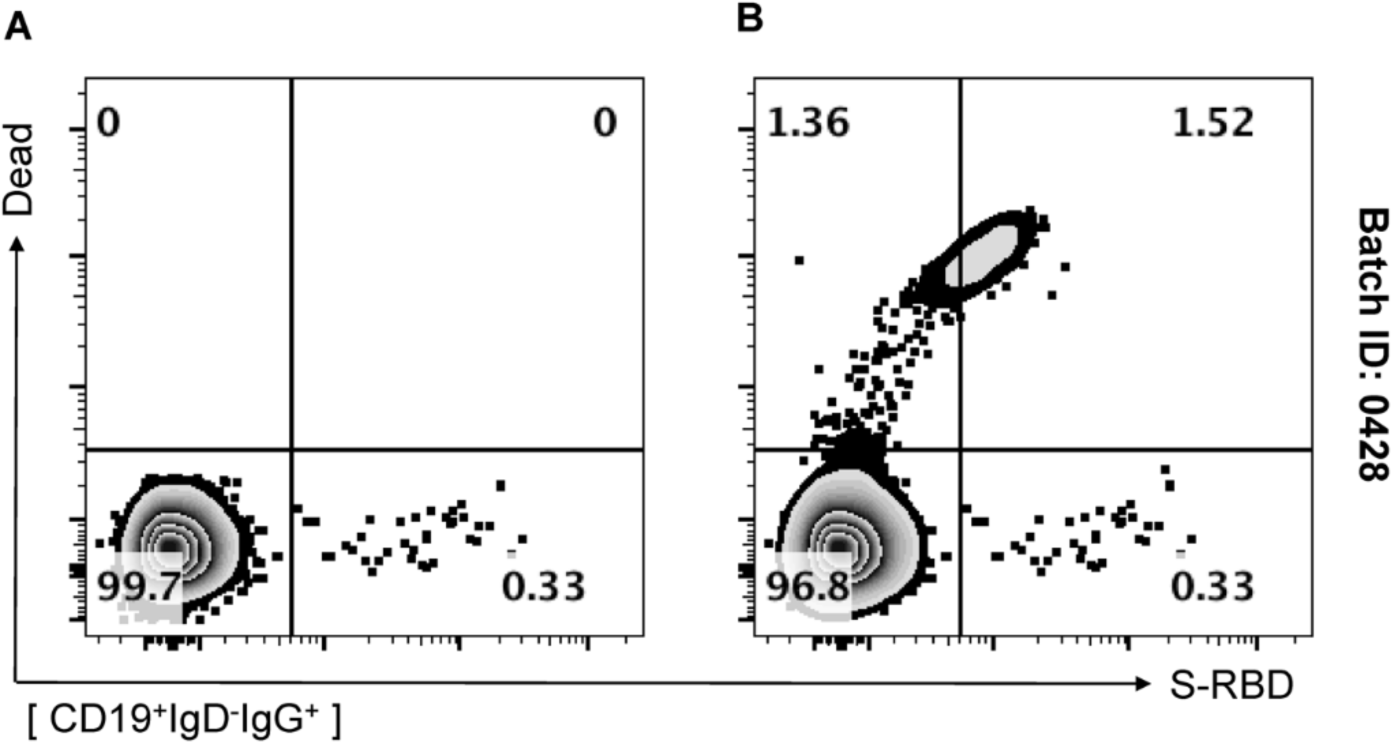
The influence of dead cells on the sorting of RBD-specific memory B cells. Gating strategy to remove dead cells: SSC-A versus FSC-A selected cell populations, then FSC-A versus FSC-H excluded doublets and FSC-H versus Dead Dye removed dead cells. Memory B cells were gated by CD19^+^IgD^−^IgG^+^ Cells (**A**), without removing dead cells in the gating strategy (**B**).

**Figure S3.**
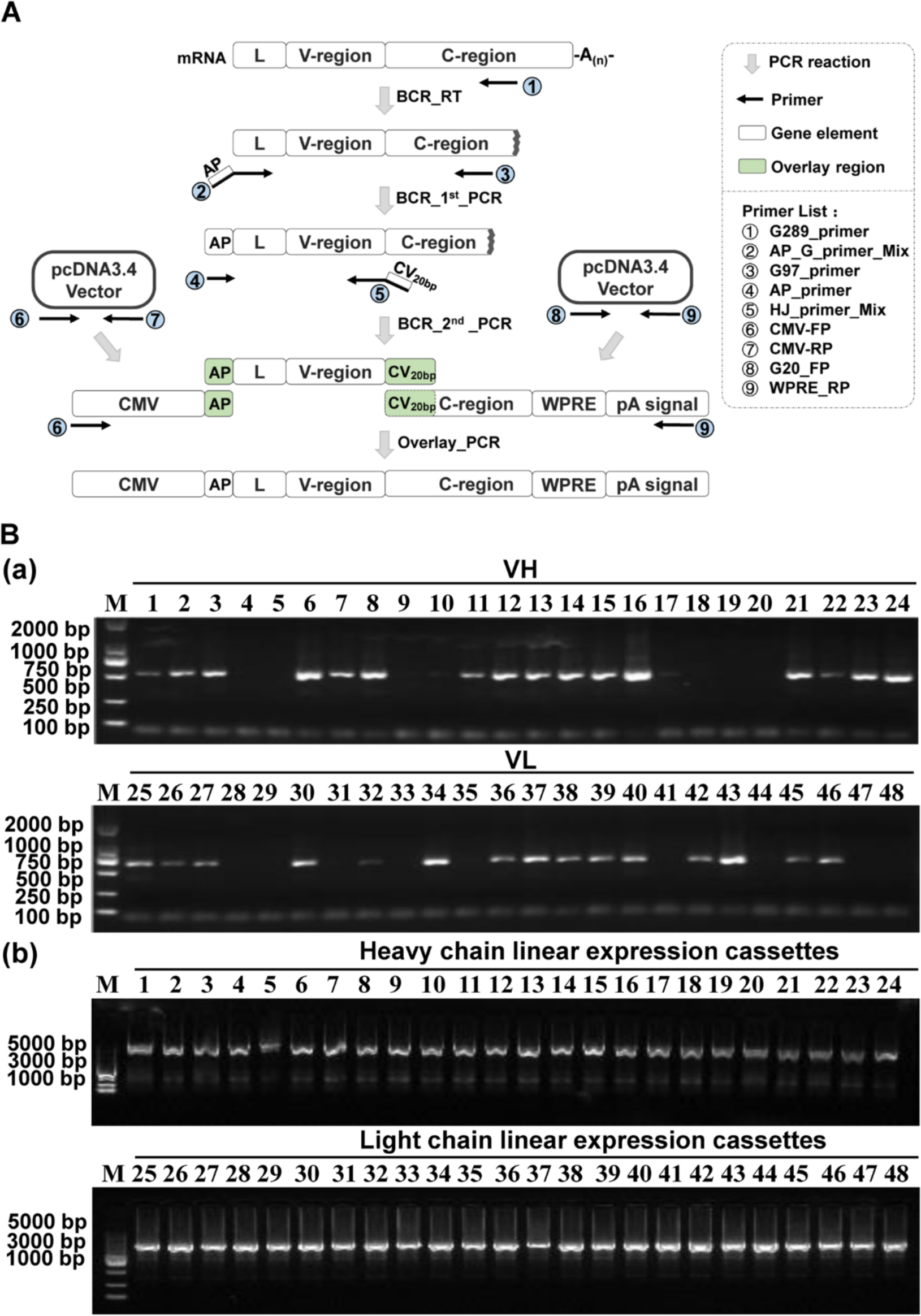
Schematic diagram of BCR RT-PCR and linear expression cassettes construction. **A.** Schema depicting workflow of the constructed linear expression cassettes. PCR amplified the variable region genes in single B cells. The BCR cDNAs was obtained from RBD-specific memory B cell by RT-PCR and the linear expression cassettes were amplified via three rounds of PCR. A primary PCR utilized gene-specific primers at both the 5′ and 3′ ends. The 5′ oligonucleotides bound the leader sequence (L). The 3′ reverse primer was connected with heavy or light constant regions. In the secondary PCR, a 5′ forward primer annealed to an "adapter", which was encoded at the 5′ end of the first PCR product, were used in combination with a 3′ reverse primer annealing to the J gene of Ab variable region. The secondary oligonucleotides provided 20 base-pair overlap regions: at the 5′ end with human cytomegalovirus (CMV) promoter fragment, and at the 3’ end with a heavy or light chain constant region fragment containing a polyadenylation sequence. Then, in a tertiary PCR, the DNAs of variable region, the CMV promoter fragment, and the constant region fragments were combined and amplified to produce two separate linear expression cassettes. **B.** The amplified products from BCR cDNAs were electrophoresed and stained with ethidium bromide. **(a)** Agarose gel of variable region BCR genes. Lane "M", 2 kb DNA ladder, Lane 1–24, heavy chain variable region and Lane 25–48, light chain variable region. **(b)** Agarose gel of linear expression cassettes. Lane "M", 5 kb DNA ladder, Lane 1–24, the linear expression cassettes of heavy chains and Lane 25–48, the linear expression cassettes of light chains.

**Figure S4.**
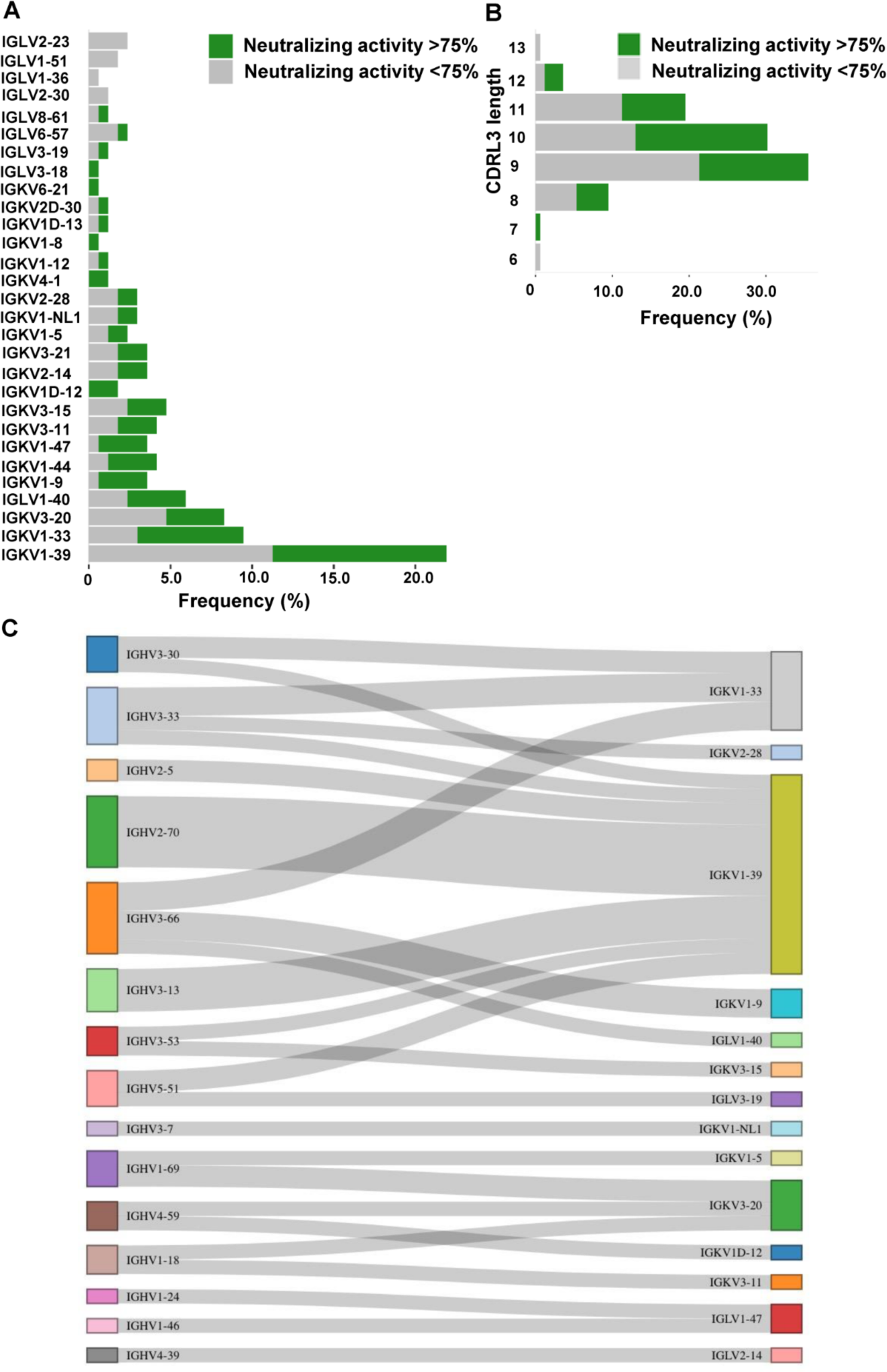
Usage and pairing of heavy and light chains for all specific antibodies. **A.** Frequencies of variable light chain gene (VL) clusters for neutralizing (activity > 75%) and potential non-neutralizing (activity < 75%) antibodies. V gene segments were ranked by frequencies of neutralizing Abs. **B.** Frequencies of various CDRL3 length of potential neutralizing and non-neutralizing antibodies. **C.** Clonal expanded heavy and light clusters were paired and highlighted in different colors.

**Figure S5.**
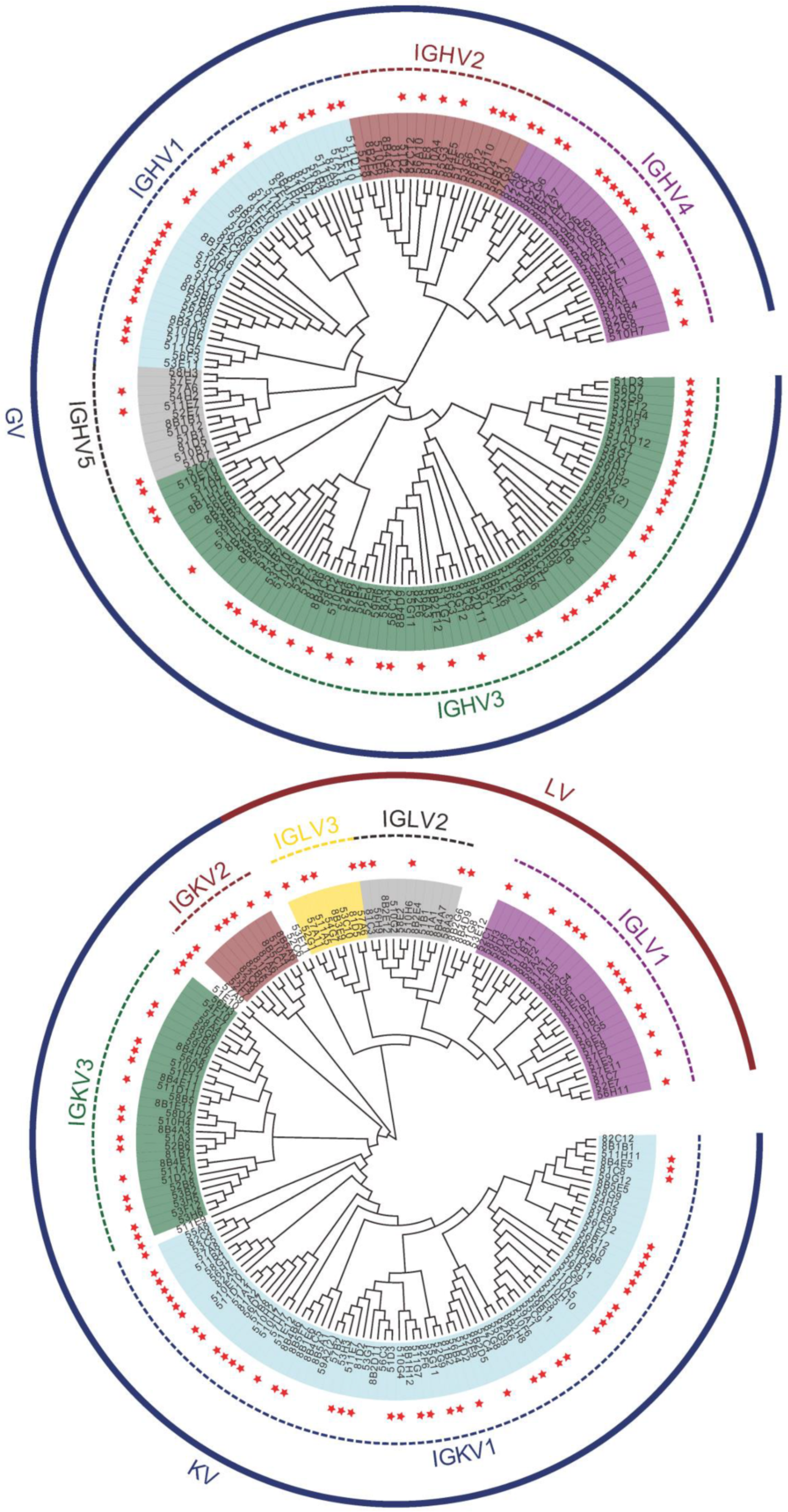
Phylogenetic analysis of VH (up) and VL (down) genes for RBD-binding antibodies. Clonal expanded VH and VL clusters were paired and highlighted in various colors. The red stars represented individual neutralizing antibodies. Branch lengths were drawn to scale so that sequence relatedness could be readily assessed.

**Table S1.**
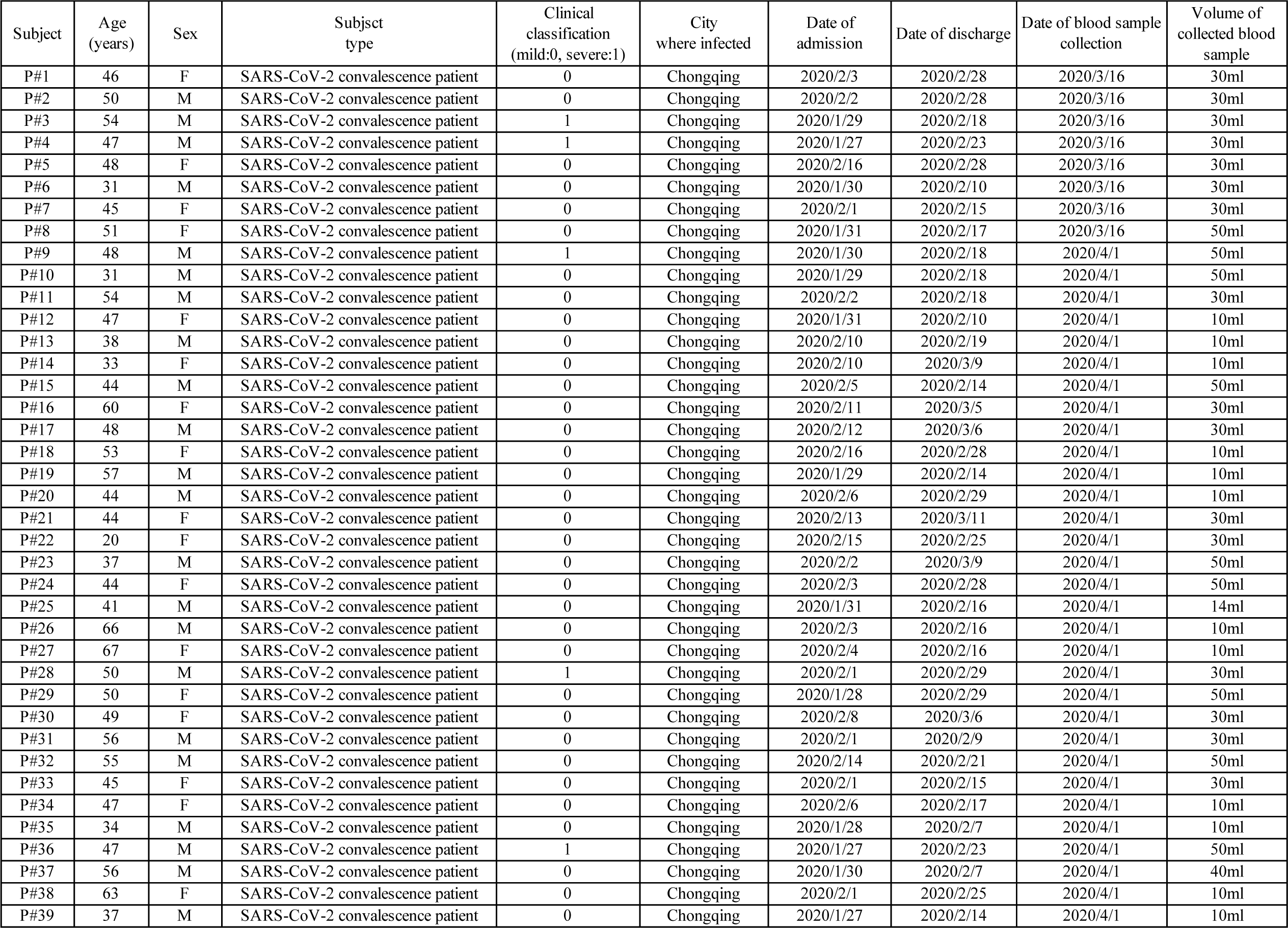
Patient information

| Subject | Age (years) | Sex | Subsct type | Clinical classification (mild:0, severe:1) | City where infected | Date of admission | Date of discharge | Date of blood sample collection | Volume of collected blood sample |
| --- | --- | --- | --- | --- | --- | --- | --- | --- | --- |
| P#1 | 46 | F | SARS-CoV-2 convalescence patient | 0 | Chongqing | 2020/2/3 | 2020/2/28 | 2020/3/16 | 30ml |
| P#2 | 50 | M | SARS-CoV-2 convalescence patient | 0 | Chongqing | 2020/2/2 | 2020/2/28 | 2020/3/16 | 30ml |
| P#3 | 54 | M | SARS-CoV-2 convalescence patient | 1 | Chongqing | 2020/1/29 | 2020/2/18 | 2020/3/16 | 30ml |
| P#4 | 47 | M | SARS-CoV-2 convalescence patient | 1 | Chongqing | 2020/1/27 | 2020/2/23 | 2020/3/16 | 30ml |
| P#5 | 48 | F | SARS-CoV-2 convalescence patient | 0 | Chongqing | 2020/2/16 | 2020/2/28 | 2020/3/16 | 30ml |
| P#6 | 31 | M | SARS-CoV-2 convalescence patient | 0 | Chongqing | 2020/1/30 | 2020/2/10 | 2020/3/16 | 30ml |
| P#7 | 45 | F | SARS-CoV-2 convalescence patient | 0 | Chongqing | 2020/2/1 | 2020/2/15 | 2020/3/16 | 30ml |
| P#8 | 51 | F | SARS-CoV-2 convalescence patient | 0 | Chongqing | 2020/1/31 | 2020/2/17 | 2020/3/16 | 50ml |
| P#9 | 48 | M | SARS-CoV-2 convalescence patient | 1 | Chongqing | 2020/1/30 | 2020/2/18 | 2020/4/1 | 50ml |
| P#10 | 31 | M | SARS-CoV-2 convalescence patient | 0 | Chongqing | 2020/1/29 | 2020/2/18 | 2020/4/1 | 50ml |
| P#11 | 54 | M | SARS-CoV-2 convalescence patient | 0 | Chongqing | 2020/2/2 | 2020/2/18 | 2020/4/1 | 30ml |
| P#12 | 47 | F | SARS-CoV-2 convalescence patient | 0 | Chongqing | 2020/1/31 | 2020/2/10 | 2020/4/1 | 10ml |
| P#13 | 38 | M | SARS-CoV-2 convalescence patient | 0 | Chongqing | 2020/2/10 | 2020/2/19 | 2020/4/1 | 10ml |
| P#14 | 33 | F | SARS-CoV-2 convalescence patient | 0 | Chongqing | 2020/2/10 | 2020/3/9 | 2020/4/1 | 10ml |
| P#15 | 44 | M | SARS-CoV-2 convalescence patient | 0 | Chongqing | 2020/2/5 | 2020/2/14 | 2020/4/1 | 50ml |
| P#16 | 60 | F | SARS-CoV-2 convalescence patient | 0 | Chongqing | 2020/2/11 | 2020/3/5 | 2020/4/1 | 30ml |
| P#17 | 48 | M | SARS-CoV-2 convalescence patient | 0 | Chongqing | 2020/2/12 | 2020/3/6 | 2020/4/1 | 30ml |
| P#18 | 53 | F | SARS-CoV-2 convalescence patient | 0 | Chongqing | 2020/2/16 | 2020/2/28 | 2020/4/1 | 10ml |
| P#19 | 57 | M | SARS-CoV-2 convalescence patient | 0 | Chongqing | 2020/1/29 | 2020/2/14 | 2020/4/1 | 10ml |
| P#20 | 44 | M | SARS-CoV-2 convalescence patient | 0 | Chongqing | 2020/2/6 | 2020/2/29 | 2020/4/1 | 10ml |
| P#21 | 44 | F | SARS-CoV-2 convalescence patient | 0 | Chongqing | 2020/2/13 | 2020/3/11 | 2020/4/1 | 30ml |
| P#22 | 20 | F | SARS-CoV-2 convalescence patient | 0 | Chongqing | 2020/2/15 | 2020/2/25 | 2020/4/1 | 30ml |
| P#23 | 37 | M | SARS-CoV-2 convalescence patient | 0 | Chongqing | 2020/2/2 | 2020/3/9 | 2020/4/1 | 50ml |
| P#24 | 44 | F | SARS-CoV-2 convalescence patient | 0 | Chongqing | 2020/2/3 | 2020/2/28 | 2020/4/1 | 50ml |
| P#25 | 41 | M | SARS-CoV-2 convalescence patient | 0 | Chongqing | 2020/1/31 | 2020/2/16 | 2020/4/1 | 14ml |
| P#26 | 66 | M | SARS-CoV-2 convalescence patient | 0 | Chongqing | 2020/2/3 | 2020/2/16 | 2020/4/1 | 10ml |
| P#27 | 67 | F | SARS-CoV-2 convalescence patient | 0 | Chongqing | 2020/2/4 | 2020/2/16 | 2020/4/1 | 10ml |
| P#28 | 50 | M | SARS-CoV-2 convalescence patient | 1 | Chongqing | 2020/2/1 | 2020/2/29 | 2020/4/1 | 30ml |
| P#29 | 50 | F | SARS-CoV-2 convalescence patient | 0 | Chongqing | 2020/1/28 | 2020/2/29 | 2020/4/1 | 50ml |
| P#30 | 49 | F | SARS-CoV-2 convalescence patient | 0 | Chongqing | 2020/2/8 | 2020/3/6 | 2020/4/1 | 30ml |
| P#31 | 56 | M | SARS-CoV-2 convalescence patient | 0 | Chongqing | 2020/2/1 | 2020/2/9 | 2020/4/1 | 30ml |
| P#32 | 55 | M | SARS-CoV-2 convalescence patient | 0 | Chongqing | 2020/2/14 | 2020/2/21 | 2020/4/1 | 50ml |
| P#33 | 45 | F | SARS-CoV-2 convalescence patient | 0 | Chongqing | 2020/2/1 | 2020/2/15 | 2020/4/1 | 30ml |
| P#34 | 47 | F | SARS-CoV-2 convalescence patient | 0 | Chongqing | 2020/2/6 | 2020/2/17 | 2020/4/1 | 10ml |
| P#35 | 34 | M | SARS-CoV-2 convalescence patient | 0 | Chongqing | 2020/1/28 | 2020/2/7 | 2020/4/1 | 10ml |
| P#36 | 47 | M | SARS-CoV-2 convalescence patient | 1 | Chongqing | 2020/1/27 | 2020/2/23 | 2020/4/1 | 50ml |
| P#37 | 56 | M | SARS-CoV-2 convalescence patient | 0 | Chongqing | 2020/1/30 | 2020/2/7 | 2020/4/1 | 40ml |
| P#38 | 63 | F | SARS-CoV-2 convalescence patient | 0 | Chongqing | 2020/2/1 | 2020/2/25 | 2020/4/1 | 10ml |
| P#39 | 37 | M | SARS-CoV-2 convalescence patient | 0 | Chongqing | 2020/1/27 | 2020/2/14 | 2020/4/1 | 10ml |

**Table S2.** Three batches of S-RBD specific B memory cell sorting

| Batch | Sample count | Subject | Volume of blood used for sorting(ml) | Total volume of blood(ml) | Number of sorted 96 wells plates | Number of Paired mAbs after BCR RT-PCR | Number of Specific mAbs (note-01) | Number of Neutralizing mAbs (note-01) |
| --- | --- | --- | --- | --- | --- | --- | --- | --- |
| 0428 | 1 | p#10 | 10 | 70 | 2 | 72 | 29 | 12 |
|  | 2 | p#11 | 10 |  |  |  |  |  |
|  | 3 | p#29 | 10 |  |  |  |  |  |
|  | 4 | p#30 | 10 |  |  |  |  |  |
|  | 5 | p#32 | 10 |  |  |  |  |  |
|  | 6 | p#36 | 10 |  |  |  |  |  |
|  | 7 | p#37 | 10 |  |  |  |  |  |
| 0505 | 1 | p#8 | 10 | 60 | 11 | 324 | 127 | 75 |
|  | 2 | p#16 | 15 |  |  |  |  |  |
|  | 3 | p#17 | 15 |  |  |  |  |  |
|  | 4 | p#26 | 10 |  |  |  |  |  |
|  | 5 | p#27 | 10 |  |  |  |  |  |
| 0528 | 1 | p#01 | 10 | 75 | 4.5 | 101 | 42 | 9 |
|  | 2 | p#03 | 10 |  |  |  |  |  |
|  | 3 | p#04 | 10 |  |  |  |  |  |
|  | 4 | p#06 | 10 |  |  |  |  |  |
|  | 5 | p#09 | 10 |  |  |  |  |  |
|  | 6 | p#22 | 10 |  |  |  |  |  |
|  | 7 | p#25 | 5 |  |  |  |  |  |
|  | 8 | p#28 | 5 |  |  |  |  |  |
|  | 9 | p#31 | 5 |  |  |  |  |  |
| Sum: |  |  |  |  |  | 497 | 198 | 96 |

Note-01: the linear expression cassettes were transfected into 293T cells for expressing Ab proteins. The supernatants of cell cultures were collected 48hs later. the specific binding ability were assessed by ELISA. the neutralization capability were assessed by pseudovirus assay.

**Table S3.**
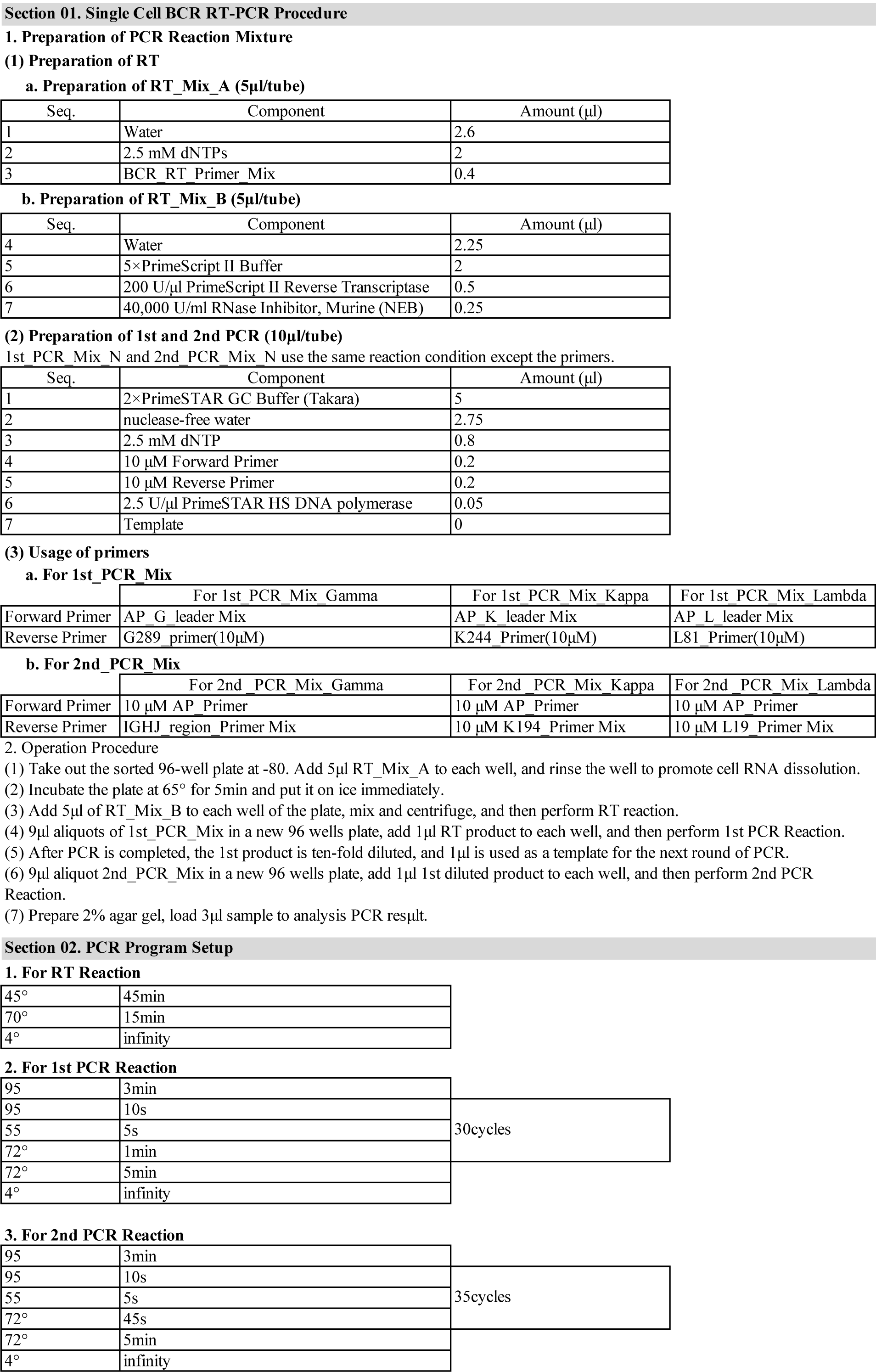
BCR RT-PCR Reaction mixture and PCR Program setup

**Table S4.**
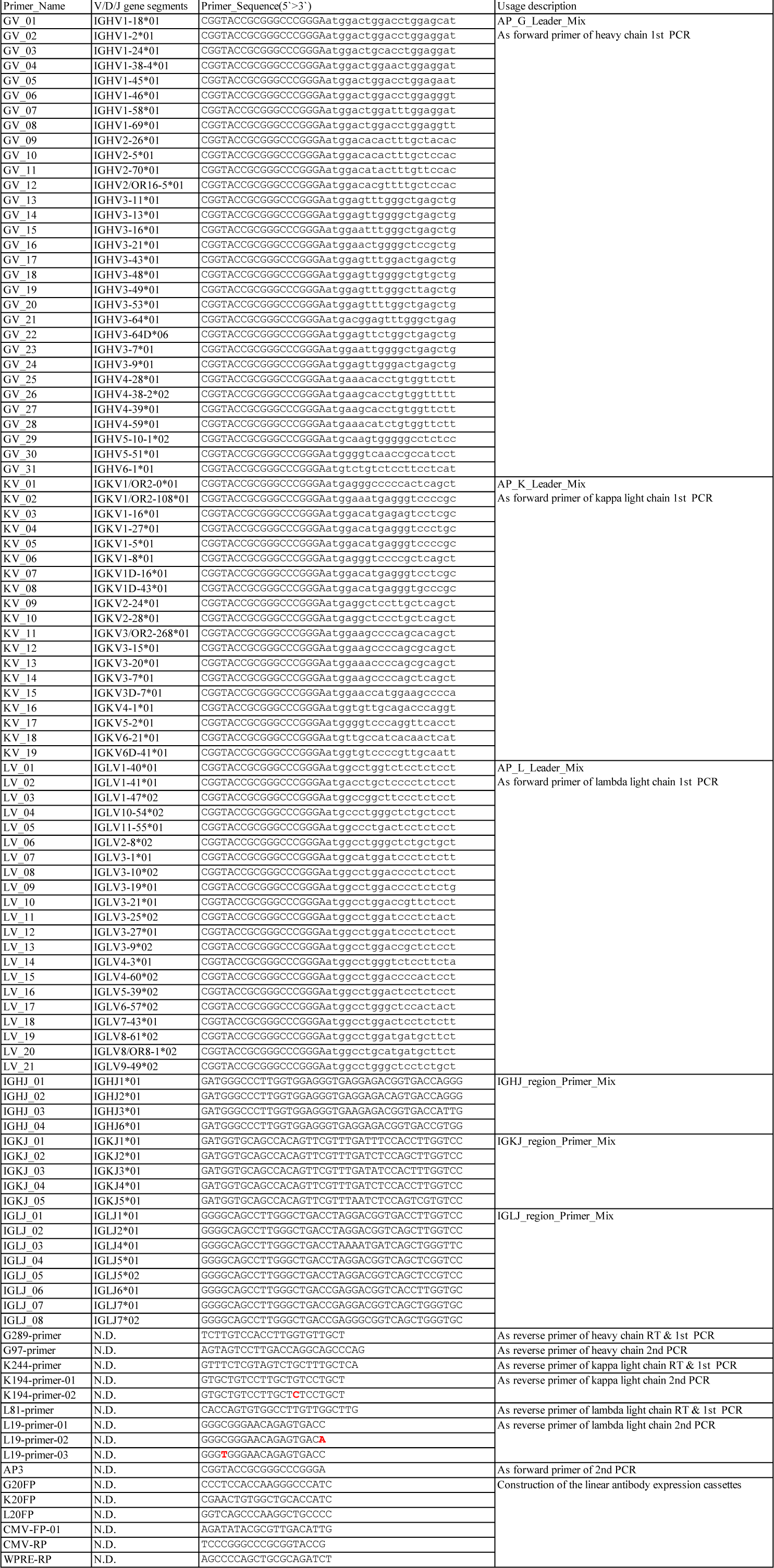
Primers List of BCR RT-PCR

**Table S5.** Preparation of BCR Cloning primers

|  |
| --- |
| **Step 01. Dissolve primer powder in water** |
| the leader primers and J-region primers are dissolved to 100uM<br>the other primers are dissolved to 10μM |
| **Step 02. Preparation of BCR_RT_Primer_Mix (each 2μM)** |
| take 100μl G289_primer(10μM), K244_primer(10μM), L81 primer(10μM), respectively, then mix according to 1:1:1. |
| **Step 03. Preparation of AP_Leader_Mix (each 2μM)** |
| (1) For AP_G_Leader_Mix: Add 380μl water to a 1.5ml centrifuge tube, and take 20μl each of the 31 GV_N primers, and the final volume is 1000μl;<br>(2) For AP_K_Leader_Mix: Add 620μl water to a 1.5ml centrifuge tube, and take 20μl each of the 19 KV_N primers, and the final volume is 1000μl;<br>(3) For AP_L_Leader_Mix: Add 580μl water to a 1.5ml centrifuge tube, and take 20μl each of the 21 LV_N primers, and the final volume is 1000μl; |
| **Step 04. Preparation of IGHJ_region_Primer_Mix (each 2μM)** |
| Add 920μl water to a 1.5ml centrifuge tube, and take 20μl each of the 4 IGHJ_N primers, and the |
| **Step 05. Preparation of K194_Primer_Mix** |
| take 100μl each of the following 2 primers, then mix:<br>K194-primer-01 (10μM)<br>K194-primer-02 (10μM) |
| **Step 06. Preparation of L19_Primer_Mix** |
| take 100μl each of the following 3 primers, then mix:<br>L19-primer-01 (10μM)<br>L19-primer-02 (10μM)<br>L19-primer-03 (10μM) |

**Table S6.**
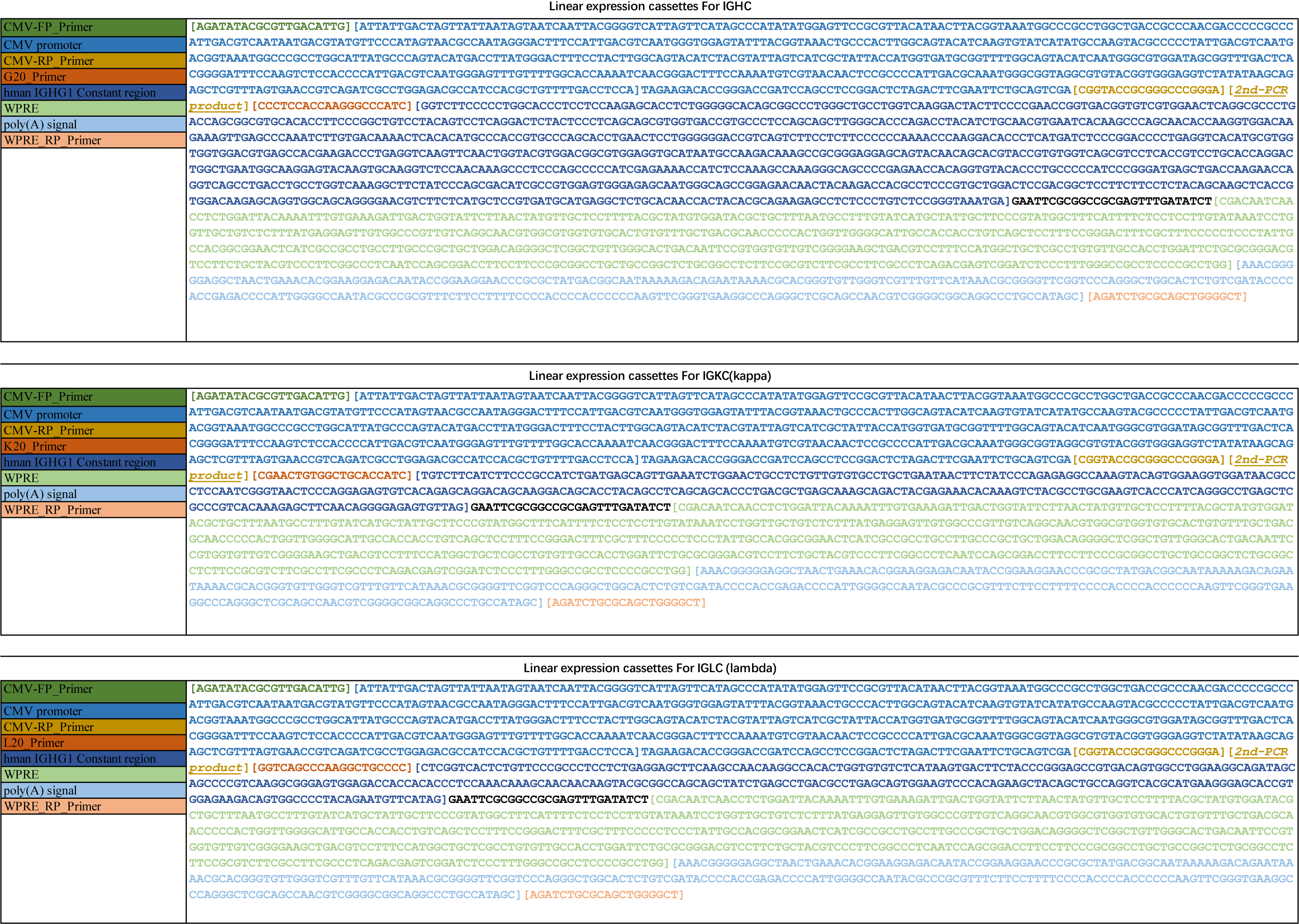
Annotation of linear antibody expression cassettes

**Table S7.**
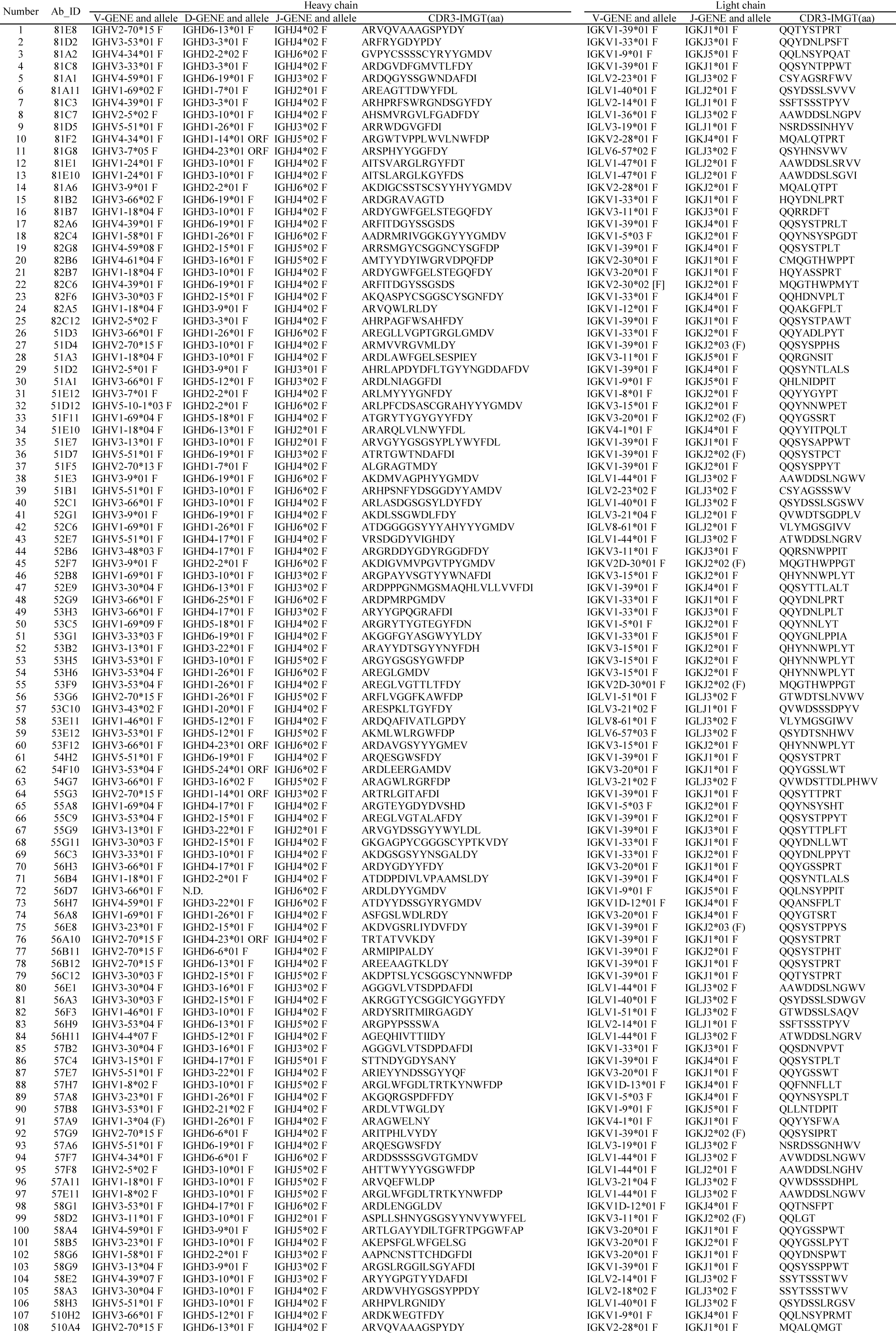
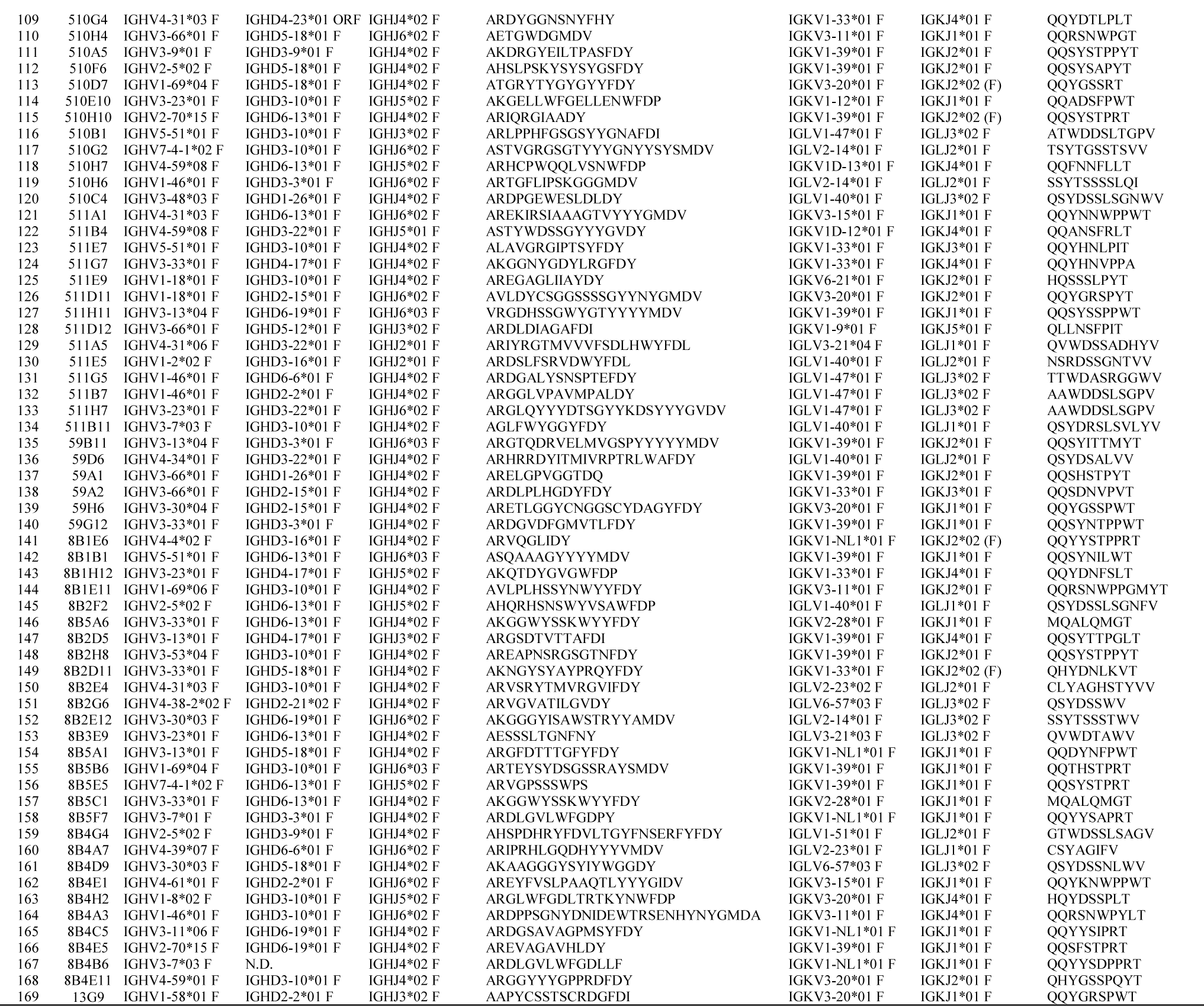
Gene family analysis of monoclonal antibodies

